# Covering soybean leaves with cellulose nanofiber changes leaf surface hydrophobicity and confers resistance against *Phakopsora pachyrhizi*

**DOI:** 10.1101/2020.08.26.267807

**Authors:** Haruka Saito, Yuji Yamashita, Nanami Sakata, Takako Ishiga, Nanami Shiraishi, Giyu Usuki, Viet Tru Nguyen, Eiji Yamamura, Yasuhiro Ishiga

**Author notes:** For correspondence: Yasuhiro Ishiga, Address: Faculty of Life and Environmental Sciences, University of Tsukuba, 1-1-1 Tennodai, Tsukuba, Ibaraki 305-8572, Japan.

## Abstract

Asian soybean rust (ASR) caused by *Phakopsora pachyrhizi*, an obligate biotrophic fungal pathogen, is the most devastating soybean production disease worldwide. Currently, timely fungicide application is the only means to control ASR in the field. We investigated cellulose nanofiber (CNF) application on ASR disease management. CNF-treated leaves showed reduced lesion number after *P. pachyrhizi* inoculation compared to control leaves, indicating that covering soybean leaves with CNF confers *P. pachyrhizi* resistance. We also demonstrated that formation of *P. pachyrhizi* pre-infection structures including germ-tubes and appressoria, and also gene expression related to these formations, such as *chitin synthases* (*CHSs*), were significantly suppressed in CNF-treated soybean leaves compared to control leaves. Moreover, contact angle measurement revealed that CNF converts soybean leaf surface properties from hydrophobic to hydrophilic. These results suggest that CNF can change soybean leaf surface hydrophobicity, conferring resistance against *P. pachyrhizi*, based on the reduced expression of *CHSs*, as well as reduced formation of pre-infection structures. This is the first study to investigate CNF application to control field disease.

## Introduction

Diseases in important crop plants have a significant negative impact on agricultural productivity. For example, Asian soybean rust (ASR) caused by *Phakopsora pachyrhizi*, an obligate biotrophic fungal pathogen, is the most devastating soybean production disease worldwide, with an estimated crop yield loss of up to 90%. ASR has impacted the South American economy in recent years. Yorinori et al. (2005) reported that the losses caused by ASR were ∼2 billion US dollars in Brazil alone in 2003. Although most rust fungi have a high host specificity, the *P. pachyrhizi* host range is broad and can infect diverse leguminous plant leaves in the field (Slaminko et al., 2008). The infection process starts when urediniospores germinate to produce a single germ-tube with an appressorium. Unlike cereal rust fungi that penetrates through stomata (Bolton et al., 2008), *P. pachyrhizi* directly penetrates into host plant epidermal cells by an appressorial peg. After penetration, *P. pachyrhizi* extends the infection hyphae and forms haustoria (feeding structures) in the mesophyll cells 24 to 48 hours after infection (Goellner et al., 2010). Five to eight days after infection, *P. pachyrhizi* then produces urediniospores by asexual reproduction (Goellner et al., 2010). Urediniospores can be dispersed by wind and germinate on other host plants.

There are several ASR control methods for soybean protection against *P. pachyrhizi*, including chemical control by fungicide application, growing ASR resistant soybean cultivars, and employing cultivation practices. Synthetic fungicides are the primary ASR disease control method. However, fungicide use can cause many problems such as environmental impacts (Maltby et al., 2009), increased production costs (Godoy et al., 2015), and the emergence of fungicide-resistant pathogens (Langenbach et al., 2016; Klosowski et al., 2018). Another major and effective control method is breeding or engineering of ASR resistant soybean cultivars. Analysis of soybean accessions disclosed six dominant *R* genes conferring resistance to a particular *P. pachyrhizi* race, and these loci were referred to as the *Rpp* 1–6 genes (Bromfield and Hartwig, 1980; McLean and Byth, 1980; Hartwig, 1986; Garcia et al., 2008; Li et al., 2012). However, none of the soybean accessions in the world show resistance to all *P. pachyrhizi* races (Monteros et al., 2007). Due to the limited resistance available in soybean cultivars, heterologous expression of resistance genes from other plant species in soybean has been investigated as an alternative source of ASR resistance. Kawashima et al. (2016) reported that soybean plants expressing *CcRpp1* (*Cajanus cajan* Resistance against *Phakopsora pachyrhizi 1*) from pigeon pea (*Cajanus cajan*) showed full resistance against *P. pachyrhizi*. Conversely, identifying resistance traits from non-host plant species has become an intelligent approach. Uppalapati et al. (2012) screened *Medicago truncatula Tnt1* mutant lines and identified an *inhibitor of rust germ tube differentiation 1* (*irg1*) mutant with reduced formation of pre-infection structures, including germ-tubes and appressoria. They demonstrated that the loss of abaxial epicuticular wax accumulation resulting in reduced surface hydrophobicity inhibited formation of pre-infection structures on the *irg1* mutant (Uppalapati et al., 2012). Furthermore, Ishiga et al. (2013) reported that gene expression related to pre-infection structure formation was activated on the hydrophobic surface of the *M. truncatula* wild-type, but not on the *irg1* mutant, based on *P. pachyrhizi* transcriptome analysis, suggesting that leaf surface hydrophobicity can trigger gene expression related to formation of pre-infection structures. Based on these previous studies, we hypothesized that modification of leaf surface hydrophobicity might be a useful strategy to confer resistance against *P. pachyrhizi*.

Cellulose is an organic polysaccharide consisting of a β-1,4 linked glucopyranose skeleton. Cellulose is an important structural component of plant primary cell walls and is essential in maintaining the plant structural phase. Due to the positive properties, cellulose has been investigated as an application in different research and development fields including energy, environmental, water, and biomedical related fields (Mondal, 2017). Cellulose nanofiber (CNF), which can be derived from cellulose, is one of the most abundant and renewable biomasses in nature (Abe et al., 2007). Because CNF exhibits properties such as low weight, high aspect ratio, high strength, high stiffness, and large surface area, CNF potentially has wide areas of application. There are several CNF isolation methods, e.g. acid hydrolysis, enzymatic hydrolysis, and mechanical processes. The aqueous counter collision (ACC) method can make it possible to cleave interfacial interactions among cellulose molecules without any chemical modification (Kondo et al., 2014). Both hydrophobic and hydrophilic sites co-exist in a cellulose molecule resulting in amphiphilic properties when CNF is derived from the ACC method. Kose et al. (2011) reported that coating with CNF derived from the ACC method could switch surface hydrophilic and hydrophobic properties, depending on substrate characteristics. They demonstrated that coating a filter paper and polyethylene with CNF changed the surface property into hydrophobic and hydrophilic, respectively (Kose et al., 2011). To investigate the potential application of CNF in agriculture, we examined whether coating with CNF protected soybean plants against *P. pachyrhizi*. We show that a specific CNF property can change soybean leaf surface hydrophobicity, resulting in reduced formation of pre-infection structures associated with reduced *P. pachyrhizi* infection.

## Materials & Methods

### Plant growth conditions, pathogen inoculation assay, and CNF treatment

Susceptible soybean cultivar seeds (*Glycine max* cv. Enrei) were germinated in a growth chamber at 25°C/20°C with 16-hrs-light/8-hrs-dark cycle (100-150 μ mol m^−2^ s^−1^) for 3 to 4 weeks.

An isolate of the ASR pathogen *P. pachyrhizi* T1-2 (Yamaoka et al., 2014) was maintained on soybean leaves. Fresh urediniospores were collected and suspended in distilled water with 0.001% Tween 20. The 3-week-old soybean plants were spray-inoculated with 1 x 10^5^ spores/ml using a hand sprayer for uniform spore deposition. The inoculated plants were maintained in a chamber for 24 hours with 90% to 95% humidity at 23°C and 0-hrs-light/24-hrs-dark cycle. The plants were then transferred to a growth chamber (22°C/20°C with 16 hrs-light/8 hrs-dark cycle) and incubated further to allow symptom development. To quantify ASR lesion number on CNF-treated plants, soybean leaves were spray-inoculated with *P. pachyrhizi*. At 10 days after inoculation, photographs were taken, and lesions were counted to calculate the lesion number per cm^2^. Lesions were counted from 54 random fields on three independent leaves.

Cellulose nanofiber (CNF, marketed as nanoforest®) supplied through the courtesy of Chuetsu Pulp & Paper (Takaoka, Japan) was used. CNF suspension was adjusted to a concentration of 0.1% including 0.02% Tween 20 (FUJIFILM, Tokyo, Japan) before treatment. Both adaxial and abaxial sides of soybean leaves were spray-treated with 0.1% CNF till runoff and then the treated soybean plants were dried at room temperature for 3 to 4 hours before inoculation. Scopoletin (TCI, Tokyo, Japan) was pre-solved as 500 mM stock solutions in DMSO and diluted to 500 μM in *P. pachyrhizi* spore suspensions.

### Quantification of pre-infection structures formation

To quantify the formation of pre-infection structures including germ-tubes and appressoria on control, CNF-, and scopletin-treated plants, soybean leaves were spray-inoculated with *P. pachyrhizi* 1 x 10^5^ spores/ml. At 6 hours after inoculation, the leaves were observed with an Olympus BX51 fluorescence microscope after Calcofluor White (Sigma-Aldrich, St. Louis, USA) staining and photographed. The germ-tubes forming differentiated appressoria were counted as appressoria. The differentiated germ-tubes without appressoria that grew on the leaf surface were also counted from at least 100 urediniosopres on three independent leaves.

The formation of pre-infection structures on borosilicate glass slides and polyethylene tape with or without CNF treatment was quantified after dropping *P. pachyrhizi* spores (2.0 x 10□/ml). Six hours after inoculation, pre-infection structures were observed with a Nikon ECLIPSE 80i phase contrast microscope. The germ-tubes forming differentiated appressoria were counted as appressoria. The differentiated germ-tubes without appressoria that grew on the leaf surface were also counted from at least 500 urediniosopres on three independent leaves.

### Contact angle measurement on soybean leaves and polyethylene tapes

The surface hydrophobicity on the CNF-treated leaves, borosilicate glass slides, and polyethylene tapes were investigated based on contact angle measurement using an automatic contact angle meter DM-31 (Kyowa Interface Science, Niiza, Japan). The contact angle was measured by dropping 2 µl of water from a syringe attached to the DM-31 automatic contact angle meter. The contact angle was measured on the adaxial and abaxial leaf surfaces, and polyethylene tapes with or without 0.1% CNF treatments. The contact angle was analyzed using the multi-functional integrated analysis software FAMAS (Kyowa Interface Science).

### RNA-spray-induced gene silencing of CHSs

Double-stranded RNA (dsRNA) of *GFP* and *CHS* were synthesized using the *in vitro* Transcription T7 Kit (Takara, Ohtsu, Japan). Briefly, we designed 3 primer sets to amplify *P. pachyrhizi CHS5-1* fragments (Supplementary Fig. S1 and Table S1). After RT-PCR amplification, fragments were purified and used as templates for *in vitro* transcription. The products of RNA transcripts were confirmed by gel electrophoresis (Supplementary Fig. S1) and quantified by NanoDrop (Thermo Fisher Scientific, Waltham, USA). We equally mixed 3 fragments and used for SIGS assay on polyethylene tape. The formation of pre-infection structures and expression levels of *CHSs* were quantified after dropping 1 x 10□/ml of P. pachyrhizi spores containing 10 ng/ml dsRNA polyethylene tape. Six hours after inoculation, pre-infection structures were observed with a Nikon ECLIPSE 80i phase contrast microscope.

### Real-time quantitative RT-PCR analyses

For urediniospores attachment assay, 4-week-old soybean leaves covered with or without 0.1% CNF were spray-inoculated with *P. pachyrhizi* 1 x 10^5^ spores/ml. The inoculated leaves were immediately fixed, and total RNA was extracted from the leaf areas and purified using RNAiso Plus (TaKaRa, Otsu, Japan). To investigate the SIGS efficacy, expression levels of *CHSs* were quantified after dropping 1 x 10□/ml of *P. pachyrhizi* spores containing 10 ng/ml dsRNA on polyethylene tape. Six hours after inoculation, total RNA was extracted from the inoculated leaf areas and purified using RNAiso Plus. To investigate the gene expression profiles of *P. pachyrhizi CHSs* during infection, 4-week-old soybean leaves were spray-inoculated with *P. pachyrhizi* 1 x 10^5^ spores/ml and incubated in darkness overnight, and then transferred to a growth chamber (22°C/20°C with a 16-h-light/8-h-dark cycle). At 2, 4, 6, 12, and 24 hours after inoculation, total RNA was extracted from the inoculated leaf areas and purified using RNAiso Plus. For gene expression profiles of *P. pachyrhizi CHSs* and soybean defense-related genes, 4-week-old soybean leaves covered with or without 0.1% CNF were spray-inoculated with *P. pachyrhizi* 1 x 10^5^ spores/ml and incubated in darkness overnight, and then transferred to a growth chamber (22°C/20°C with a 16-h-light/8-h-dark cycle). At 6, 12, and 24 hours after inoculation, total RNA was extracted from the inoculated leaf areas and purified using RNAiso Plus according to the manufacture’s protocol.

Two micrograms of total RNA were treated with gDNA Remover (TOYOBO, Osaka, Japan) to eliminate genomic DNA, and the DNase-treated RNA was reverse transcribed using the ReverTra Ace qPCR RT Master Mix (TOYOBO). The cDNA (1:10) was then used for RT-qPCR using the primers shown in Supplementary Table S1 with THUNDERBIRD SYBR qPCR Mix (TOYOBO) on a Thermal Cycler Dice Real Time System (TaKaRa). *P. pachyrhizi Ubiquitin 5* (*PpUBQ5*) and soybean *Ubiquitin* (*GmUBQ3*) were used to compare urediniospores attachment on soybean leaves. *P. pachyrhizi Elongation factor 1α* (*PpEF1α*) and *Ubiquitin 5* (*PpUBQ5*) were used to normalize *P. pachyrhizi* gene expression. Soybean *GmEF1α* and *GmUBQ3* were used as internal controls to normalize soybean gene expression.

## Results

### Covering soybean leaves with CNF confers resistance against *P. pachyrhizi*

To investigate the potential application of CNF in agriculture, especially disease resistance against pathogens, we first treated soybean leaves with CNF. Four hours after spraying with 0.1% CNF, we challenged soybean leaves with *P. pachyrhizi* and observed lesion formation including uredinia at 10 days after inoculation. CNF-treated leaves showed reduced lesion area compared to control leaves (Fig. 1A). CNF-treated leaves showed significantly reduced lesion number compared to control leaves (Fig. 1B). These results indicate that covering soybean leaves with CNF confers resistance against *P. pachyrhizi*. Next, we investigated urediniospores attachment on control and CNF-treated leaves by quantifying the relative ratio of *ubiquitin* gene transcripts in soybean and *P. pachyrhizi*. As shown in Fig. 1C, we found no significant difference in the relative ratio of *ubiquitin* transcripts between control and CNF-treated leaves, indicating that urediniospores were uniformly sprayed on control and CNF-treated leaves.

**Figure 1.**
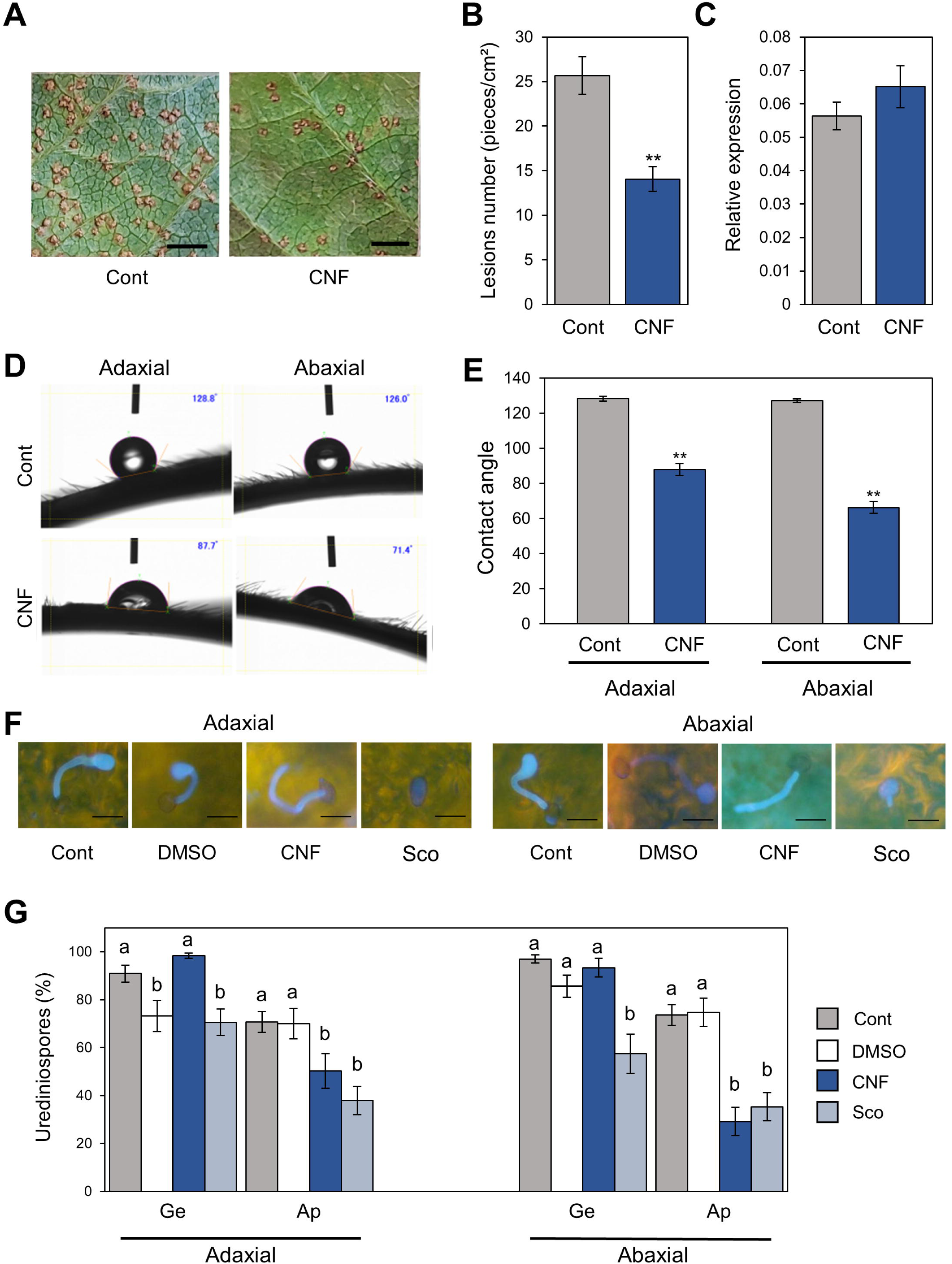
*P. pachyrhizi* lesion formation, pre-infection structures formation, and hydrophobicity on CNF-treated soybean leaves. **(A)** Lesions resulting from *P. pachyrhizi* infection on the abaxial leaf surface of control, and leaves covered with 0.1% cellulose nanofiber derived (CNF). Soybean plants were spray-inoculated with *P. pachyrhizi* (1 x 10^5^ spores/ml), and photographs were taken 10 days after inoculation. Bars indicate 0.2 cm. **(B)** Lesion numbers resulting from *P. pachyrhizi* infection on the abaxial leaf surface of control, and leaves covered with 0.1% cellulose nanofiber (CNF). Soybean plants were spray-inoculated with *P. pachyrhizi* (1 x 10^5^ spores/ml), and lesion numbers were counted to calculate lesion number per cm^2^. Vertical bars indicate the standard error of the means (*n* = 54). Asterisks indicate a significant difference between control and CNF-treatments in a *t* test (** *p* < 0.01). **(C)** Urediniospore attachment quantification on the leaf surface of control and leaves covered with 0.1% cellulose nanofiber derived (CNF). Soybean plants were spray-inoculated with *P. pachyrhizi* (1 x 10^5^ spores/ml) and immediately total RNAs including soybean and *P. pachyrhizi* were purified. Relative expression of soybean *Ubiquitin 3* (*GmUBQ3*) and *P. pachyrhizi Ubiquitin 5* (*PpUBQ5*) were evaluated using RT-qPCR. Vertical bars indicate the standard error of the means (*n* = 4). **(D)** Reduction of contact angle and hydrophobicity on cellulose nanofiber derived (CNF)-treated soybean leaves. Contact angles of water droplets on the adaxial and abaxial leaf surface of control, and leaves covered with 0.1% cellulose nanofiber (CNF). Contact angles were evaluated as described in the Methods. **(E)** Reduction of contact angle and hydrophobicity on cellulose nanofiber derived (CNF)-treated soybean leaves. Contact angles of water droplets on the adaxial and abaxial leaf surface of control, and leaves covered with 0.1% cellulose nanofiber (CNF). Contact angles were evaluated as described in the Methods. Vertical bars indicate the standard error of the means (*n* = 60). Asterisks indicate a significant difference between control and CNF-treatments in a *t* test (** *p* < 0.01). **(F)** Suppression of *P. pachyrhizi* pre-infection structures on cellulose nanofiber (CNF)-treated soybean leaves. *P. pachyrhizi* pre-infection structure formation on the adaxial and abaxial surfaces of control, and leaves covered with 0.1% cellulose nanofiber (CNF), treated with 0.1% DMSO and 500 μM scopoletin. Soybean plants were spray-inoculated with *P. pachyrhizi* (1 x 10^5^ spores/ml). The pre-infection structures were stained with Calcofluor White and photographs were taken 6 hours after inoculation. Bars indicate 50 μm. **(G)** Suppression of *P. pachyrhizi* pre-infection structures on cellulose nanofiber (CNF)-treated soybean leaves. *P. pachyrhizi* pre-infection structure formation on the adaxial and abaxial surfaces of control, and leaves covered with 0.1% cellulose nanofiber (CNF), treated with 0.1% DMSO and 500 μM scopoletin. Soybean plants were spray-inoculated with *P. pachyrhizi* (1 x 10^5^ spores/ml). The photographs were taken 6 hours after inoculation, and the percentage of germinated (Ge) urediniospores and differentiated germ-tubes with appressoria (Ap) were evaluated as described in the Methods. Vertical bars indicate the standard error of the means (*n* = 21). Significant differences (*p* < 0.05) are indicated by different letters based on a Tukey’s honestly significant difference (HSD) test.

### CNF converts leaf surface properties from hydrophobic to hydrophilic

CNF has amphipathic properties, and thus can convert material surface properties from hydrophobic to hydrophilic, and *vice versa* (Kose et al., 2011). To confirm whether CNF-treatments can convert soybean leaf surface properties from hydrophobic to hydrophilic, we quantified the differences in surface hydrophobicity by measuring the contact angle at the interface of a liquid (water) drop with the leaf surface. A greater contact angle (>90°) is indicative of poor wetting or hydrophobicity. Interestingly, significant differences in the contact angle were observed between control and CNF-treated adaxial leaf surfaces (Fig. 2D). The adaxial leaf surface of control leaves exhibited an average contact angle of 128°, whereas CNF-treated leaves showed a dramatic decrease in the contact angle (around 90°), which is indicative of a hydrophilic surface (Fig. 2E). Similarly, significant differences in the contact angle were observed between control and CNF-treated abaxial leaf surfaces (Fig. 2D). The abaxial leaf surface of control leaves exhibited an average contact angle of 127°, whereas CNF-treated leaves showed a dramatic decrease in contact angle (around 70°; Fig. 2E). These results clearly indicate that CNF-treatments can convert leaf surface properties from hydrophobic to hydrophilic.

**Figure 2.**
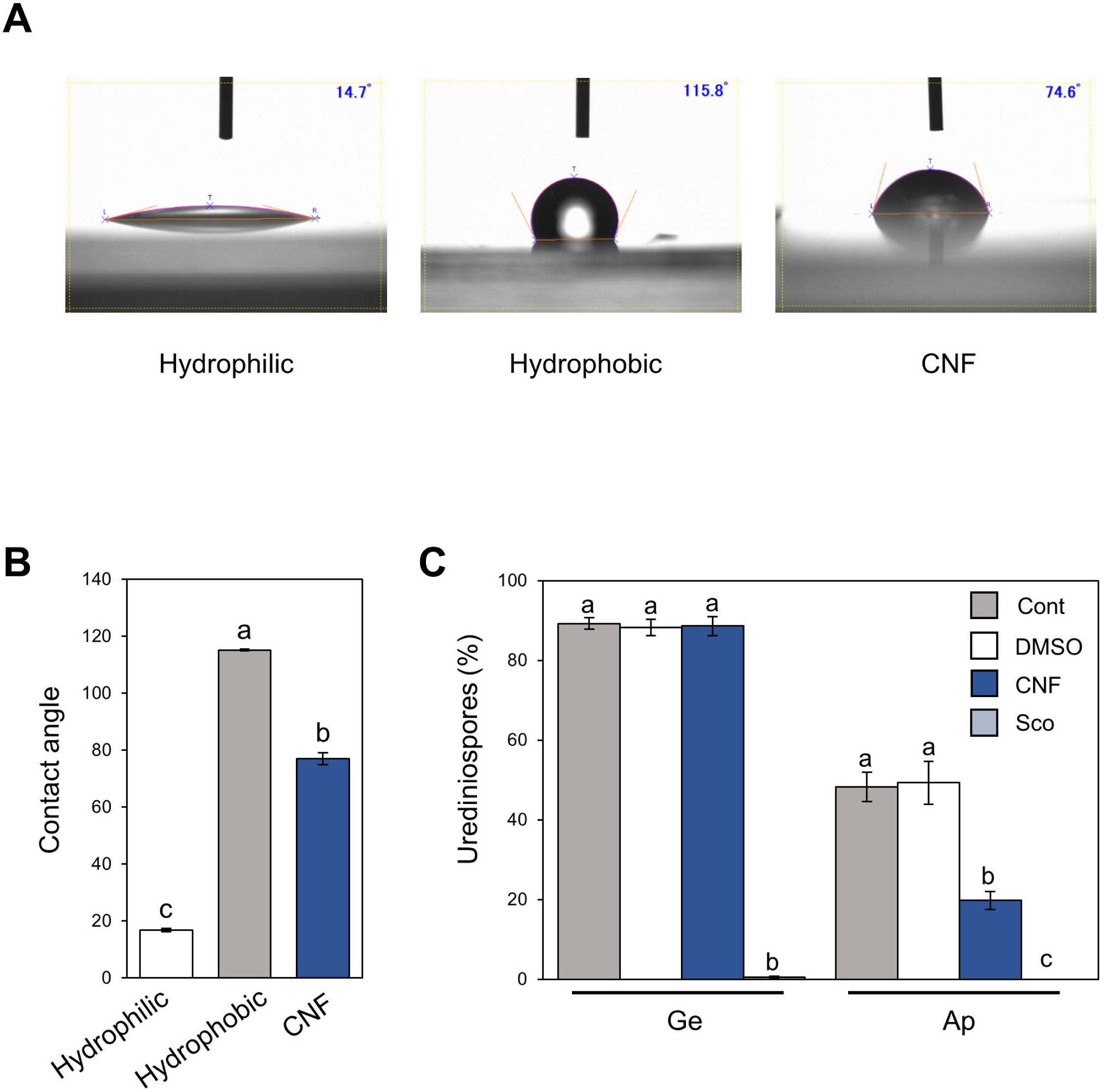
*P. pachyrhizi* pre-infection structures formation and hydrophobicity on polyethylene surfaces covered with CNF. **(A)** Hydrophobicity on borosilicate glass slide (hydrophilic), and polyethylene tape (hydrophobic) covered with or without 0.1% cellulose nanofiber (CNF). Contact angles were evaluated as described in Methods. **(B)** Hydrophobicity on borosilicate glass slide (hydrophilic), and polyethylene tape (hydrophobic) covered with or without 0.1% cellulose nanofiber (CNF). Contact angles were evaluated as described in Methods. Significant differences (*p* < 0.05) are indicated by different letters based on a Tukey’s honestly significant difference (HSD) test. **(C)** Suppression of *P. pachyrhizi* pre-infection structures on cellulose nanofiber (CNF)-treated polyethylene tape. *P. pachyrhizi* pre-infection structure formation on polyethylene tape covered with or without 0.1% cellulose nanofiber (CNF), treated with 0.1% DMSO and 500 μM scopoletin. Polyethylene tapes were spray-inoculated with *P. pachyrhizi* (1 x 10^5^ spores/ml). The photographs were taken 6 h after inoculation and the percentage of germinated (Ge) urediniospores and differentiated germ-tubes with appressoria (Ap) were evaluated as described in the Methods. Vertical bars indicate the standard error of the means (*n* = 19 ∼ 28). Significant differences (*p* < 0.05) are indicated by different letters based on a Tukey’s honestly significant difference (HSD) test.

### Covering soybean leaves with CNF suppresses formation of *P. pachyrhizi* pre-infection structures

Since CNF-treatments suppressed the lesion number, we next investigated the formation of pre-infection structures including germ-tubes and appressoria on CNF-treated leaves. In control leaves, around 90% of urediniospores germinated, and ∼75% formed appressoria on adaxial and abaxial leaves (Fig. 1F and Fig. 1G). In CNF-treated leaves, around 90% of urediniospores germinated, and interestingly ∼50% and ∼30% of them formed appressoria on adaxial and abaxial leaves, respectively (Fig. 1F and Fig. iG). We also investigated the scopoletin application effect, since scopoletin is known to protect soybean from soybean rust by suppressing the formation of pre-infection structures (Beyer et al., 2019). Consistent with a previous study, in scopoletin-treated leaves, ∼70% and ∼60% of urediniospores germinated, and ∼40% and ∼30% of them formed appressoria on adaxial and abaxial leaves, respectively (Fig. 1F and Fig. 1G). These results suggest that covering soybean leaves with CNF suppresses formation of pre-infection structures, including germ-tubes and appressoria.

### Hydrophobicity with CNF suppresses formation of *P. pachyrhizi* pre-infection structures

Since CNF-treatments converted leaf surface properties from hydrophobic to hydrophilic, and suppressed the formation of pre-infection structures, we next investigated the effect of CNF treatment on hydrophobic polyethylene tape. The hydrophilic borosilicate glass slide exhibited an average contact angle of 16.8°, whereas the hydrophobic polyethylene tape showed an average contact angle of 115.1° (Fig. 2A and 2B). Interestingly, CNF-treated polyethylene tape showed a dramatic decrease in contact angle (around 90°), which is indicative of a hydrophilic surface (Fig. 2A and 2B). On control polyethylene tape, around 90% of urediniospores germinated, and ∼50% formed appressoria on hydrophobic surfaces (Fig. 2C). On CNF-treated polyethylene tape, around 90% of urediniospores germinated, and interestingly ∼20% of them formed appressoria (Fig. 2C). Scopoletin suppressed urediniospore germination (Fig. 2C). These results suggest that covering hydrophobic surfaces with CNF suppresses formation of pre-infection structures, including germ-tubes and appressoria, which resulted from conversion of surface properties from hydrophobic to hydrophilic.

### *P. pachyrhizi* chitin synthases are required for formation of pre-infection structures

Ishiga et al. (2013) reported that gene expression related to formation of pre-infection structures was induced on the hydrophobic surface based on *P. pachyrhizi* transcriptome analysis. Chitin synthases (CHSs) are key enzymes in the biosynthesis of the fungal cell wall structural component, chitin. Since Ishiga et al. (2013) demonstrated that *P. pachyrhizi CHS* expression was induced on the hydrophobic leaf surface, we next tested the expression profiles of *P. pachyrhizi CHS* genes in soybean leaves. Except for *CHS2-1* and *CHS3-3*, all *CHS* gene transcripts were significantly induced within 2 hours after soybean leaf inoculation (Fig. 3A and Supplementary Fig. 1A), suggesting CHSs may be involved in the formation of pre-infection structures, including germ-tubes and appressoria.

**Figure 3.**
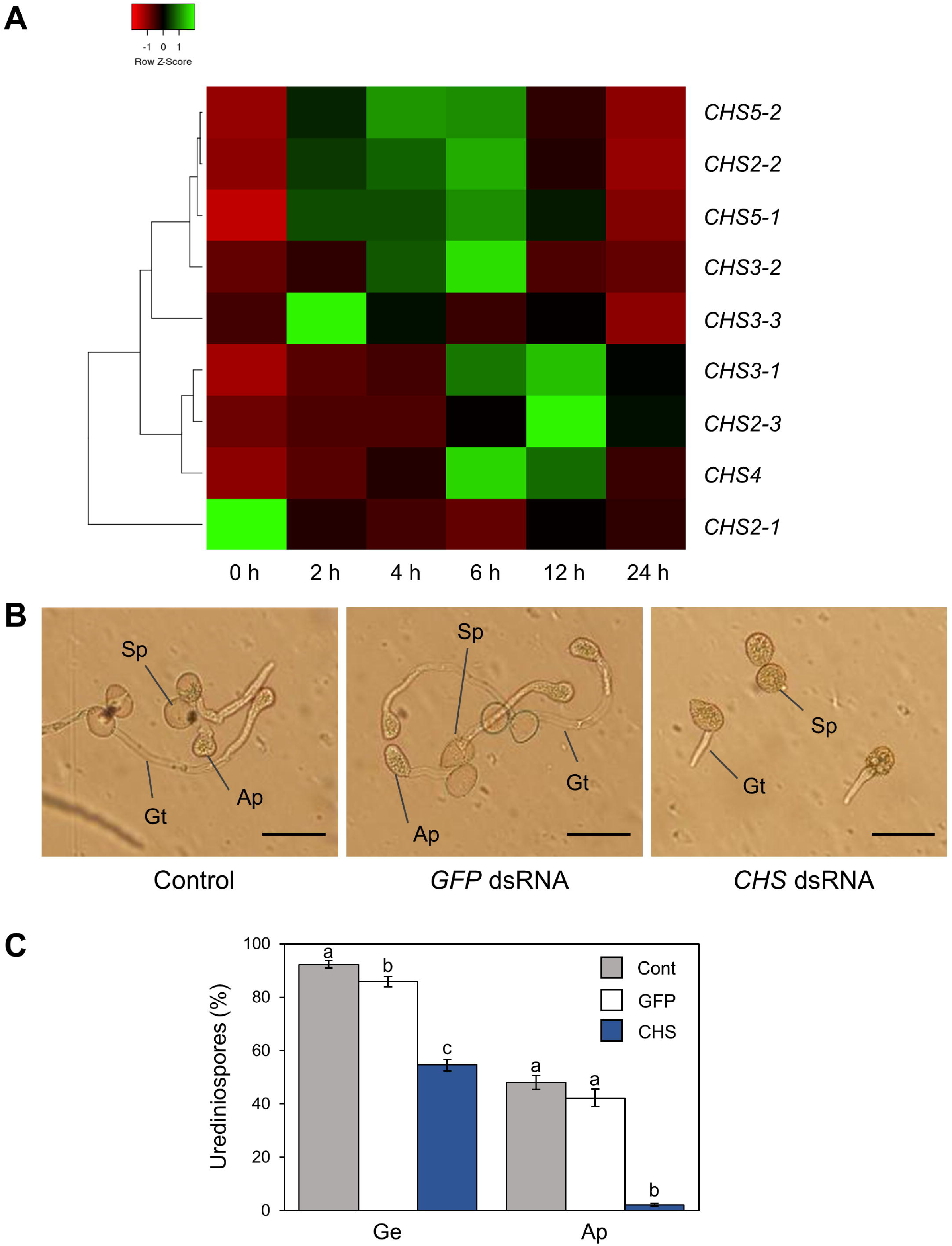
Gene expression profiles and functional analysis of *P. pachyrhizi chitin synthase* genes. **(A)** The heatmap created from gene expression profiles of *P. pachyrhizi chitin synthases*, including *CHS2-1*, *CHS2-2*, *CHS2-3*, *CHS3-1*, *CHS3-2*, *CHS3-3*, *CHS4*, *CHS5-1*, and *CHS5-2* on soybean leaves. Soybean plants were spray-inoculated with *P. pachyrhizi* (1 x 10^5^ spores/ml). Total RNAs including soybean and *P. pachyrhizi* was purified at 0, 2, 4, 6, 12, and 24 hours after inoculation, and expression profiles were evaluated using RT-qPCR. *P. pachyrhizi Elongation factor* and *Ubiquitin 5* were used to normalize the samples. Expression profiles were visualized as a heatmap using Heatmapper (Babicki et al., 2016). **(B)** Suppression of *P. pachyrhizi* pre-infection structures by *chitin synthase* (*CHS*) double-stranded RNA (dsRNA). *P. pachyrhizi* pre-infection structure formation was observed on polyethylene tapes treated with *GFP* dsRNA and *CHS* dsRNA. Polyethylene tapes were spray-inoculated with *P. pachyrhizi* (1 x 10^5^ spores/ml). The photographs were taken 6 h after inoculation. Bars indicate 50 μm. **(C)** Suppression of *P. pachyrhizi* pre-infection structures by *chitin synthase* (*CHS*) double-stranded RNA (dsRNA). *P. pachyrhizi* pre-infection structure formation on polyethylene tapes treated with *GFP* dsRNA and *CHS* dsRNA. Polyethylene tapes were spray-inoculated with *P. pachyrhizi* (1 x 10^5^ spores/ml). The photographs were taken 6 hours after inoculation and the percentage of germinated (Ge) urediniospores and differentiated germ-tubes with appressoria (Ap) were evaluated as described in the Methods. Vertical bars indicate the standard error of the means (*n* = 46 ∼ 47). Significant differences (*p* < 0.05) are indicated by different letters based on a Tukey’s honestly significant difference (HSD) test.

To investigate *P. pachyrhizi* CHSs function on pre-infection structures formation, we performed RNA-spray-induced gene silencing (SIGS) targeting *CHS* genes. We designed dsRNA to target all *CHS* genes, and checked these gene expression levels on a hydrophobic polyethylene surface with or without *CHS* dsRNA for 6 hours. As expected, all *CHS* transcripts were significantly suppressed by treatment with *CHS* dsRNA (Supplementary Fig. S2). We next investigated the effect of *CHS* dsRNA on pre-infection structures formation. On control polyethylene tape with *GFP* dsRNA treatment, around 90% of urediniospores germinated, and ∼50% of them formed appressoria on the hydrophobic surface (Fig. 3B and 3C). Interestingly, with CHS dsRNA treatment, around ∼60% of urediniospores germinated, and interestingly less than 5% of them formed appressoria (Fig. 3B and 3C). These results clearly indicate that *P. pachyrhizi* CHSs are required for formation of pre-infection structures, including germ-tubes and appressoria.

### Soybean defense-related gene expression analysis

Nanofibers such as chitin nanofibers induce plant immune responses by activating defense-related gene expression (Egusa et al., 2015). Therefore, one could argue that the CNF-induced resistance phenotype in soybean plants may result from defense response activation, rather than from the direct effects of CNF treatments against *P. pachyrhizi*. To rule out this possibility, we investigated the expression profiles of the defense marker *PR* genes and defense-related genes, including phenylpropanoid and isoflavonoid pathways leading to phytoalexin production. Except *IFR* and *CHR*, all defense marker *PR* genes and defense-related genes were clearly induced within 6 hours of *P. pachyrhizi* inoculation, and these transcripts reached high levels at 12 hours (Fig. 4 and Supplementary Fig. S3). Interestingly, the transcript levels of defense marker *PR* genes and defense-related genes were significantly less at 6 hours on CNF-treated soybean leaves compared to control leaves, suggesting that CNF treatment does not induce *PR* and defense-related genes. These results confirmed that the resistance phenotype against *P. pachyrhizi* on CNF-treated soybean leaves is a direct effect of CNF treatment.

**Figure 4.**
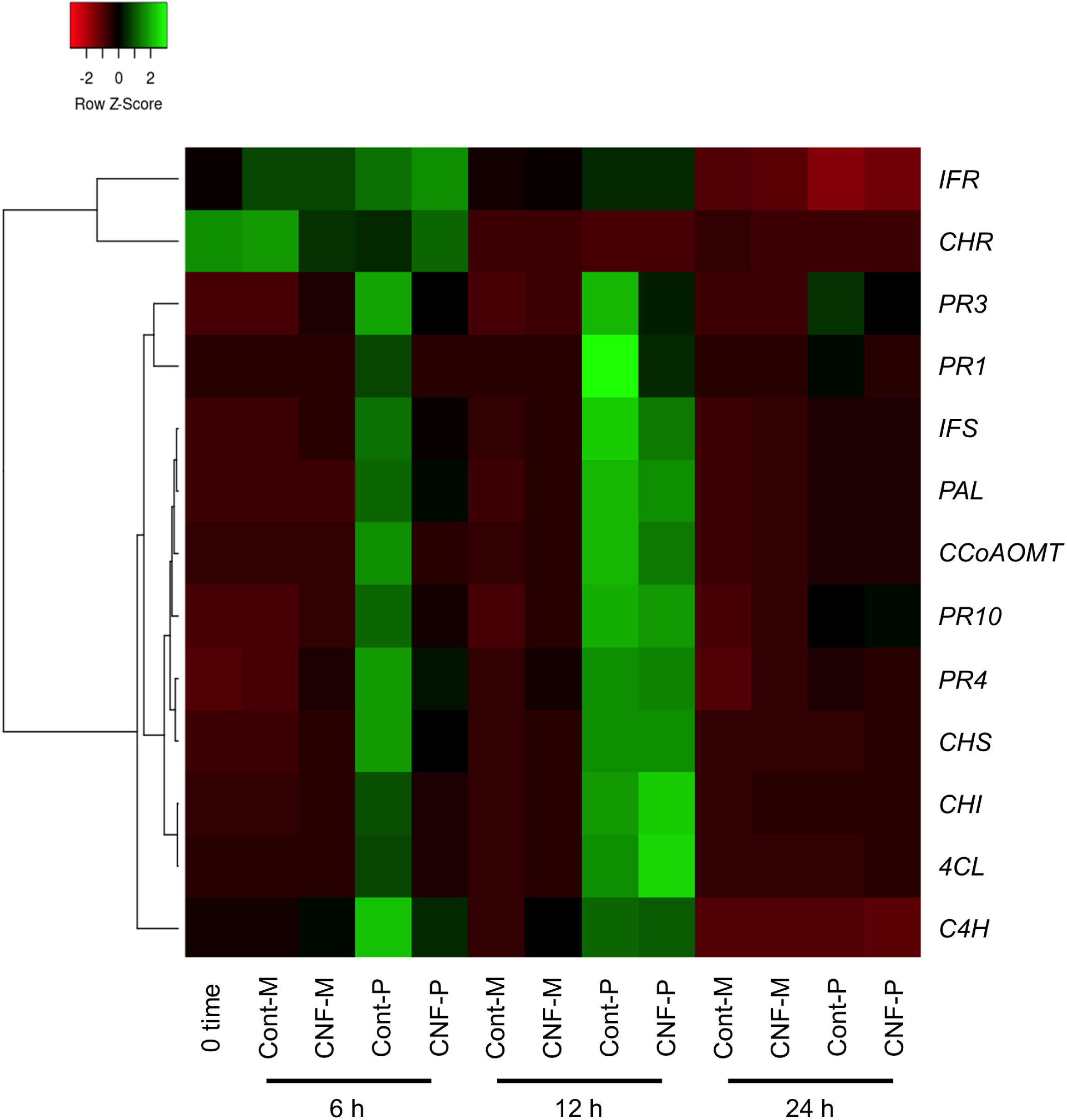
Gene expression profiles of soybean defense marker *PR* and defense-related genes in response to *P. pachyrhizi* inoculation on CNF-treated leaves. The heatmap was created from gene expression profiles of soybean defense marker *PR* and defense-related genes including *pathogenesis-related protein 1* (*PR1*), *2* (*PR2*), *3* (*PR3*), *4* (*PR4*), *10* (*PR10*), *phenylalanine ammonia-lyase* (*PAL*), *cinnamate 4-hydroxylase* (*C4H*), *4-coumarate CoA ligase* (*4CL*), *caffeoyl coenzyme A O-methyltransferase* (*CCoAOMT*), *chalcone synthase* (*CHS*), *chalcone reductase* (*CHR*), *chalcone isomerase* (*CHI*), isoflavone synthase (*IFS*), and *isoflavone reductase* (*IFR*) in response to *P. pachyrhizi* inoculation on CNF-treated leaves. Soybean plants were spray-inoculated with *P. pachyrhizi* (1 x 10^5^ spores/ml). Total RNAs including soybean and *P. pachyrhizi* were purified at 6, 12, and 24 hours after inoculation and expression profiles were evaluated using RT-qPCR. Soybean *elongation factor 1α* (*GmEF1α*) and *ubiquitin 3* (*GmUBQ3*) were used to normalize the samples. Expression profiles were visualized as a heatmap using Heatmapper (Babicki et al., 2016).

### Covering soybean leaves with CNF changes gene expression profiles related to formation of pre-infection structures

*P. pachyrhizi* CHSs are required for formation of pre-infection structures (Fig. 3). We next investigated gene expression profiles of *CHSs* in control and CNF-treated leaves at 6, 12, and 24 hours after *P. pachyrhizi* inoculation. Except *CHS2-1* and *CHS3-3*, all *CHSs* gene transcripts were clearly induced within 6 h in control soybean leaves (Fig. 5). However, the expression of these genes was clearly suppressed in CNF-treated leaves (Fig. 5A, 5C, 5D, 5E, 5F, 5H, 5I, and 5J), indicating that covering soybean leaves with CNF changes gene expression profiles of CHSs. Together, these results suggest that CNF-treatments suppress the expression of *CHSs*, resulting in reduced chitin biosynthesis activity in the *P. pachyrhizi* cell wall.

**Figure 5.**
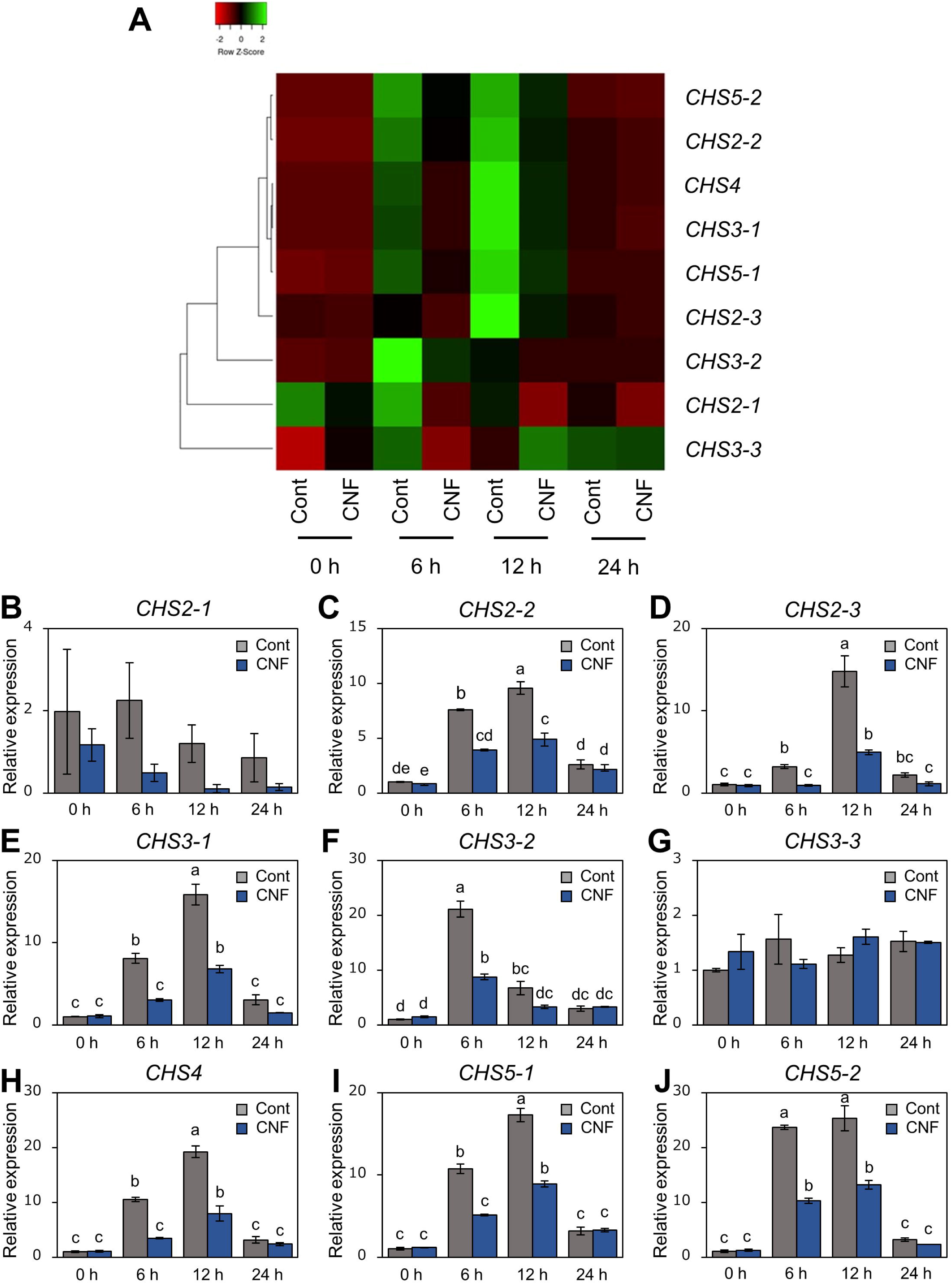
Gene expression profiles of *P. pachyrhizi chitin synthase* genes on CNF-treated soybean leaves. **(A)** The heatmap was created from gene expression profiles of *P. pachyrhizi chitin synthases*, including *CHS2-1*, *CHS2-2*, *CHS2-3*, *CHS3-1*, *CHS3-2*, *CHS3-3*, *CHS4*, *CHS5-1*, and *CHS5-2* on soybean leaves covered with or without 0.1% cellulose nanofiber (CNF). Soybean plants were spray-inoculated with *P. pachyrhizi* (1 x 10^5^ spores/ml). Total RNAs including soybean and *P. pachyrhizi* were purified at 0, 6, 12, and 24 hours after inoculation, and expression profiles were evaluated using RT-qPCR. *P. pachyrhizi elongation factor 1α* (*PpEF1α*) and *ubiquitin 5* (*PpUBQ5*) were used to normalize the samples. Expression profiles were visualized as a heatmap using Heatmapper (Babicki et al., 2016). Gene expression profiles of *P. pachyrhizi chitin synthases*, including *CHS2-1* (**B**), *CHS2-2* (**C**), *CHS2-3* (**D**), *CHS3-1* (**E**), *CHS3-2* (**F**), *CHS3-3* (**G**), *CHS4* (**H**), *CHS5-1* (**I**), and *CHS5-2* (**J**) on soybean leaves covered with or without 0.1% cellulose nanofiber (CNF). Vertical bars indicate the standard error of the means (*n* = 4). Significant differences (*p* < 0.05) are indicated by different letters based on a Tukey’s honestly significant difference (HSD) test.

**Figure 6.**
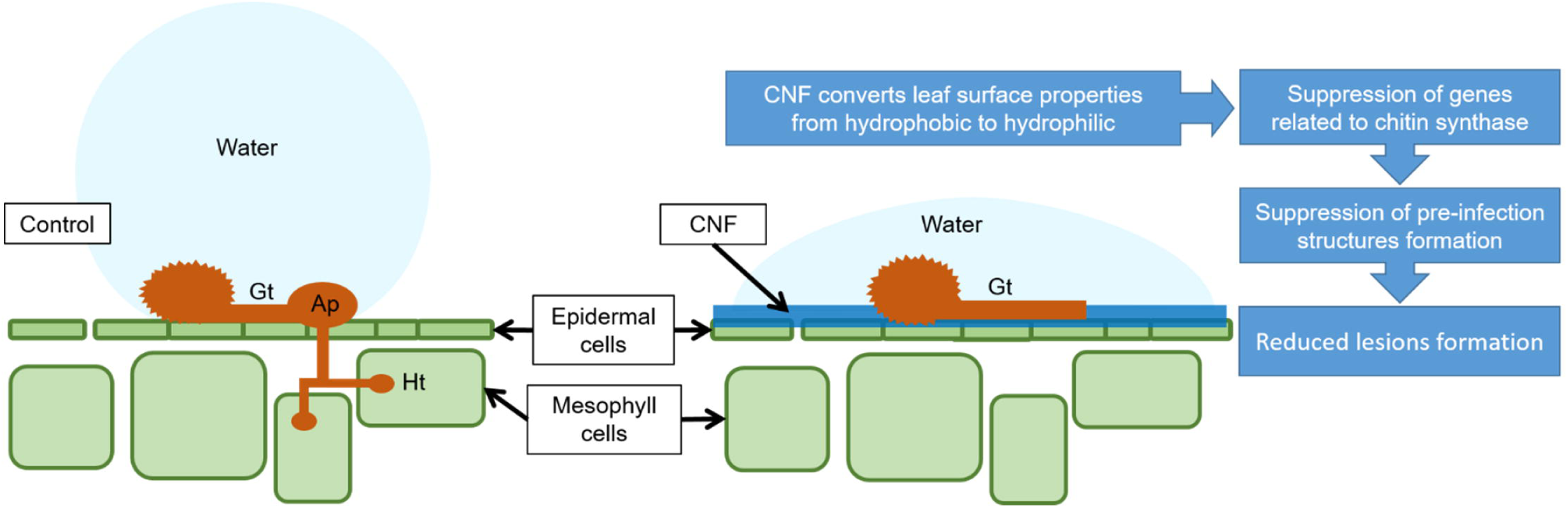
Proposed mechanism model by which CNF-treatments confer resistance against *P. pachyrhizi*. CNF-treatments convert leaf surface properties from hydrophobic to hydrophilic. The formation of pre-infection structures, and the associated gene expressions related to these formations are suppressed on CNF-treated leaves, resulting in reduced *P. pachyrhizi* infection. Gt, Ap, and Ht show germ-tubes, appressoria, and haustoria, respectively.

## Discussion

We investigated the potential application of CNF in agriculture, especially disease protection, and found that CNF-treated soybean leaves conferred resistance against the rust pathogen *P. pachyrhizi* (Fig. 1A and Fig. 1B). CNF-treatments convert soybean leaf surface properties from hydrophobic to hydrophilic (Fig. 1D and Fig. 1E), resulting in suppression of *P. pachyrhizi CHSs* genes involved in the formation of pre-infection structures, including germ-tubes and appressoria (Fig. 5) associated with reduced appressoria formation (Fig. 1F and 1G). These results provide new insights into CNF application on *P. pachyrhizi* disease management strategies.

CNF-treatments conferred soybean resistance against *P. pachyrhizi* associated with reduced lesion formation (Fig. 1A and Fig. 1B). The application of chitin nanofibers for plant protection against pathogens has been investigated. Egusa et al. (Egusa et al., 2015) reported that chitin nanofibers effectively reduced fungal and bacterial pathogen infections in *Arabidopsis thaliana* by activating plant defense responses, including reactive oxygen species (ROS) production and defense-related gene expression. Furthermore, chitin nanofiber treatment can reduce the occurrence of Fusarium wilt disease in tomato plants (Egusa et al., 2019). These results suggest that chitin nanofibers activate plant immunity, resulting in reduced pathogen infection. However, we showed no CNF elicitor activity based on defense gene expression profiles (Fig. 4). Although there is no similarity to the mechanism by which nanofibers, including cellulose and chitin, function to protect plants against pathogens, both nanofibers will be able to provide eco-friendly disease control strategies in sustainable agriculture.

Formation of pre-infection structures including germ-tubes and appressoria was significantly suppressed in CNF-treated leaves compared to control leaves (Fig. 2). Consistent with our results, Uppalapati et al. (Uppalapati et al., 2012) reported the reduced formation of pre-infection structures on a *M. truncatula irg1* mutant, in which the epicuticular waxes were completely defective and the surface property was changed to hydrophilic. These results indicate that properties such as hydrophobicity are important to form *P. pachyrhizi* pre-infection structures during early infection stages. The importance of hydrophobicity and/or epicuticular waxes on the formation of germ-tubes and appressoria has also been reported for other fungal pathogens (Mendoza-Mendoza et al., 2009; Hansjakob et al., 2010; Weidenbach et al., 2014). Further characterization of the mechanisms by which fungal pathogens recognize plant surface properties and initiate infection behavior will be needed to develop effective and sustainable disease control methods.

CNF-treatments suppressed *P. pachyrhizi CHSs* expression related to chitin formation, which are associated with reduced formation of pre-infection structures (Fig. S2, Fig. 2 and Fig. 3). CHSs are important in cell wall formation in most filamentous fungi (Takeshita et al., 2005; Lenardon et al., 2010). Treitschke et al. (2010) reported that an *Ustilago maydis CHS5* mutant Δ*msc1* showed reduced virulence associated with abnormal hyphal morphology. Madrid et al. (2003) also demonstrated that CHS5 in *Fusarium oxysporum*, a causal agent of tomato vascular wilt, has a crucial role in virulence and mediates the tomato protective response. A *F. oxysporum CHS5* mutant could not infect tomato, and exhibited abnormal morphologies such as hyphal swelling, due to changes in the cell wall properties (Madrid et al., 2003). These results suggest that *CHS5* gene deficiency or mutation causes morphological abnormalities in fungal cell wall formation, leading to virulence suppression. Together, it is tempting to speculate that suppression of *P. pachyrhizi CHS5* in CNF-treated leaves may result in changes in the cell wall properties of *P. pachyrhizi* pre-infection structures. Further characterization of CHSs, especially CHS5 based on dsRNA-mediated silencing such as SIGS and host-induced gene silencing (HIGS), in conjunction with analysis of *P. pachyrhizi* cell wall properties on CNF-treated leaves, will be necessary to understand CHSs molecular function during formation of pre-infection structures.

We demonstrated that CNF-treatments suppressed ASR caused by *P. pachyrhizi*, one of the most important soybean diseases (Fig. 1A and Fig. 1B) associated with reduced formation of pre-infection structures (Fig. 1F and Fig. 1G). Because numerous rust and filamentous fungal pathogens form pre-infection structures during early infection stages, these results imply that CNF might be an additional disease management tool to prevent crop diseases against these pathogens. However, we tested the ability of CNF to protect plants against an obligate biotrophic pathogen, but not other pathogen types, including hemibiotrophs and necrotrophs. Therefore, further characterization of CNF effects on disease suppression not only against fungal pathogens, but also against bacterial pathogens will be needed.

Our results demonstrated that SIGS targeting *P. pachyrhizi CHSs* functioned successfully in reducing pre-infection structures formation on hydrophobic polyethylene surfaces (Fig. 3B, 3C, and Supplementary Fig. 2). SIGS is a technology that promotes silencing by spraying the target dsRNA on the plant surface. Therefore, it is possible to silence a specific phytopathogen gene and protect the plant without the need for plant gene recombination (Cagliari et al., 2019; Wytinck et al., 2020). Hu et al., (2020) demonstrated that SIGS targeting *P. pachyrhizi* genes encoding an acetyl-CoA acyltransferase, a 40S ribosomal protein S16, and glycine cleavage system H protein reduced pustule numbers over 70%. SIGS against filamentous fungi threating agronomically important crops has also been studied, including head blight caused by *Fusarium graminearum* and gray mold caused by *Botrytis cinerea* (Koch et al., 2016; Wang et al., 2016; Nerva et al., 2020). Koch et al. (2016) demonstrated that SIGS targeting *Fusarium graminearum cytochrome P450* genes, which are required for fungal ergosterol biosynthesis, successfully inhibited fungal growth in barley. Although further precise studies for SIGS targeting *P. pachyrhizi* virulence genes will be needed, SIGS is a powerful tool to develop sustainable disease management strategies.

Expression profiles of soybean leaves revealed that gene transcripts related to the phenylpropanoid and isoflavonoid pathways were upregulated within 6 hours of *P. pachyrhizi* inoculation, and reached high levels at 12 hours (Fig. 4). Consistent with our results, previous studies reported that the expression of these genes was upregulated within 12 hours after *P. pachyrhizi* inoculation (Naoumkina et al., 2010; Schneider et al., 2011; Ishiga et al., 2015; Hossain et al., 2018). We demonstrated that the transcript levels of defense marker *PR* genes and defense-related genes were significantly less at 6 hours in CNF-treated soybean leaves compared to control leaves (Fig. 4). Further, appressoria formation was significantly reduced in CNF-treated leaves compared to controls (Fig. 1F and 1G). Therefore, it is tempting to speculate that reduced transcripts of defense marker *PR* genes and defense-related genes in CNF-treated leaves is the result of the decreased penetration rate associated with reduced appressoria formation.

In summary, CNF-treatments confer resistance against *P. pachyrhizi*, a causal agent of ASR. Moreover, CNF-treatments can change leaf surface hydrophobicity, resulting in CHSs gene suppression related to chitin synthase, which is associated with reduced formation of pre-infection structures including *P. pachyrhizi* germ-tubes and appressoria (Fig. 5). Since CNF is an abundant and renewable biomass in nature, CNF application for plant protection will provide a new avenue into eco-friendly and sustainable disease management.

## Figure legends

**Supplementary Figure S1. Expression profiles of *P. pachyrhizi chitin synthases* (*CHSs*), including *CHS2-1* (A), *CHS2-2* (B), *CHS2-3* (C), *CHS3-1* (D), *CHS3-2* (E), *CHS3-3* (F), *CHS4* (G), *CHS5-1* (H), and *CHS5-2* (I) on soybean leaves.** Soybean plants were spray-inoculated with *P. pachyrhizi* (1 x 10^5^ spores/ml). Total RNAs including soybean and *P. pachyrhizi* were purified at 0, 2, 4, 6, 12, and 24 hours after inoculation, and expression profiles were evaluated using RT-qPCR. *P. pachyrhizi elongation factor 1α* (*PpEF1α*) and *ubiquitin 5* (*PpUBQ5*) were used to normalize the samples. Vertical bars indicate the standard error of the means (*n* = 4). Asterisks indicate a significant difference between 0 h and each time point in a *t* test (* *p* < 0.05, ** *p* < 0.01).

**Supplementary Figure S2. Expression profiles of *P. pachyrhizi chitin synthases* (*CHSs*), including *CHS2-1* (A), *CHS2-2* (B), *CHS2-3* (C), *CHS3-1* (D), *CHS3-2* (E), *CHS3-3* (F), *CHS4* (G), *CHS5-1* (H), and *CHS5-2* (I) on polyethylene tapes treated with double-stranded RNA.** Down-regulation of *CHSs* transcripts were quantified on polyethylene tapes treated with *GFP* dsRNA and *CHS* dsRNA. Polyethylene tapes were spray-inoculated with *P. pachyrhizi* (1 x 10^5^ spores/ml). Total RNAs of *P. pachyrhizi* were purified 6 hours after inoculation, and expression profiles were evaluated using RT-qPCR. *P. pachyrhizi elongation factor 1α* (*PpEF1α*) and *ubiquitin 5* (*PpUBQ5*) were used to normalize the samples. Vertical bars indicate the standard error of the means (*n* = 4). Asterisks indicate a significant difference between 0 time and each time point in a *t* test (* *p* < 0.05, ** *p* < 0.01).

**Supplementary Figure S3. Gene expression profiles of soybean defense marker PR and defense-related genes in response to P. pachyrhizi inoculation on CNF-treated leaves.** Gene expression profiles of soybean defense marker *PR* and defense-related genes including *pathogenesis-related protein 1* (*PR1*; **A**), *2* (*PR2*; **B**), *3* (*PR3*; **C**), *4* (*PR4*; **D**), *10* (*PR10*; **E**), *phenylalanine ammonia-lyase* (*PAL*; **F**), *cinnamate 4-hydroxylase* (*C4H*; **G**), *4-coumarate CoA ligase* (*4CL*; **H**), *caffeoyl coenzyme A O-methyltransferase* (*CCoAOMT*; **H**), *chalcone synthase* (*CHS*; **I**), *chalcone reductase* (*CHR*; **J**), *chalcone isomerase* (*CHI*; **K**), *isoflavone synthase* (*IFS*; **L**), and *isoflavone reductase* (*IFR*; **M**) in response to *P. pachyrhizi* inoculation on CNF-treated leaves. Soybean plants were spray-inoculated with *P. pachyrhizi* (1 x 10^5^ spores/ml). Total RNAs including soybean and *P. pachyrhizi* were purified at 6, 12, and 24 hours after inoculation, and expression profiles were evaluated using RT-qPCR. Soybean *elongation factor 1α* (*GmEF1α*) and *ubiquitin 3* (*GmUBQ3*) were used to normalize the samples. Significant differences (*p* < 0.05) are indicated by different letters based on a Tukey’s honestly significant difference (HSD) test.

**Supplementary Figure S4**. Gene sequence and primer sets for *in vitro* transcription of *CHS5-1* double-stranded RNA

**Supplementary Table S1**. Primer sets used in this study

## References

1. Abe, K., Iwamoto, S., and Yano, H. (2007). Obtaining cellulose nanofibers with a uniform width of 15 nm from wood. Biomacromolecules 8, 3276–3278. doi: 10.1021/bm700624p

2. Babicki, S., Arndt, D., Marcu, A., Liang, Y., Grant, J.R., Maciejewski, A., and Wishart, D.S. (2016). Heatmapper: web-enabled heat mapping for all. Nucleic Acids Res. 44, W147–153. doi: 10.1093/nar/gkw419

3. Beyer, S.F., Beesley, A., Rohmann, P.F.W., Schultheiss, H., Conrath, U., and Langenbach, C.J.G. (2019). The *Arabidopsis* non-host defense-associated coumarin scopoletin protects soybean from Asian soybean rust. Plant J. 99, 397–413. doi: 10.1111/tpj.14426

4. Bolton, M.D., Kolmer, J.A., and Garvin, D.F. (2008). Wheat leaf rust caused by *Puccinia triticina*. Mol. Plant Pathol. 9, 563–575. doi: 10.1111/j.1364-3703.2008.00487.x

5. Bromfield, K.R., and Hartwig, E.E. (1980). Resistance to soybean rust and mode of inheritance. Crop Sci. 20, 254–255. doi: 10.2135/CROPSCI1980.0011183X002000020026X

6. Cagliari, D., Dias, N.P., Galdeano, D.M., Dos Santos, E., Smagghe, G., and Zotti, M.J. (2019). Management of pest insects and plant diseases by non-transformative RNAi. Front. Plant Sci. 10, 1319. doi: 10.3389/fpls.2019.01319

7. Egusa, M., Parada, R., Aklog, Y.F., Ifuku, S., and Kaminaka, H. (2019). Nanofibrillation enhances the protective effect of crab shells against Fusarium wilt disease in tomato. Int. J. Biol. Macromol. 128, 22–27. doi: 10.1016/j.ijbiomac.2019.01.088

8. Egusa, M., Matsui, H., Urakami, T., Okuda, S., Ifuku, S., Nakagami, H., and Kaminaka, H. (2015). Chitin nanofiber elucidates the elicitor activity of polymeric chitin in plants. Front. Plant Sci. 6, 1098. doi: 10.3389/fpls.2015.01098

9. Garcia, A., Calvo, E.S., de Souza Kiihl, R.A., Harada, A., Hiromoto, D.M., and Vieira, L.G. (2008). Molecular mapping of soybean rust (*Phakopsora pachyrhizi*) resistance genes: discovery of a novel locus and alleles. Theor. Appl. Genet. 117, 545–553. doi: 10.1007/s00122-008-0798-z

10. Godoy, C.V., Bueno, A.d.F., and Gazziero, D.L.P. (2015). Brazilian soybean pest management and threats to its sustainability. Outlooks on Pest Management 26, 3.

11. Goellner, K., Loehrer, M., Langenbach, C., Conrath, U., Koch, E., and Schaffrath, U. (2010). *Phakopsora pachyrhizi*, the causal agent of Asian soybean rust. Mol. Plant Pathol. 11, 169–177. doi: 10.1111/j.1364-3703.2009.00589.x

12. Hansjakob, A., Bischof, S., Bringmann, G., Riederer, M., and Hildebrandt, U. (2010). Very-long-chain aldehydes promote *in vitro* prepenetration processes of *Blumeria graminis* in a dose- and chain length-dependent manner. New Phytol. 188, 1039–1054. doi: 10.1111/j.1469-8137.2010.03419.x

13. Hartwig, E.E. (1986). Identification of a fourth major gene conferring resistance to soybean rust. Crop Sci. 26, 1135–1136. doi: 10.2135/cropsci1986.0011183X002600060010x

14. Hossain, M.Z., Ishiga, Y., Yamanaka, N., Ogiso-Tanaka, E., and Yamaoka, Y. (2018). Soybean leaves transcriptomic data dissects the phenylpropanoid pathway genes as a defence response against *Phakopsora pachyrhizi*. Plant Physiol. Biochem. 132, 424–433. doi: 10.1016/j.plaphy.2018.09.020

15. Hu, D., Chen, Z.Y., Zhang, C., and Ganiger, M. (2020). Reduction of *Phakopsora pachyrhizi* infection on soybean through host- and spray-induced gene silencing. Mol. Plant Pathol. 21, 794–807. doi: 10.1111/mpp.12931

16. Ishiga, Y., Upplapapti, S., and Mysore, K.S. (2013). Expression analysis reveals a role for hydrophobic or epicuticular wax signals in pre-penetration structure formation of *Phakopsora pachyrhizi*. Plant Signal Behav. 8, e26959. doi: 10.4161/psb.26959

17. Ishiga, Y., Uppalapati, S.R., Gill, U.S., Huhman, D., Tang, Y., and Mysore, K.S. (2015). Transcriptomic and metabolomic analyses identify a role for chlorophyll catabolism and phytoalexin during *Medicago* nonhost resistance against Asian soybean rust. Sci. Rep. 5, 13061. doi: 10.1038/srep13061

18. Kawashima, C.G., Guimarães, G.A., Nogueira, S.R., MacLean, D., Cook, D.R., Steuernagel, B., Baek, J., Bouyioukos, C., Melo, B.o.V., Tristão, G., de Oliveira, J.C., Rauscher, G., Mittal, S., Panichelli, L., Bacot, K., Johnson, E., Iyer, G., Tabor, G., Wulff, B.B., Ward, E., Rairdan, G.J., Broglie, K.E., Wu, G., van Esse, H.P., Jones, J.D., and Brommonschenkel, S.H. (2016). A pigeonpea gene confers resistance to Asian soybean rust in soybean. Nat. Biotechnol. 34, 661–665. doi: 10.1038/nbt.3554

19. Klosowski, A.C., Castellar, C., Stammler, G., and May De Mio, L.L. (2018). Fungicide sensitivity and monocyclic parameters related to the *Phakopsora pachyrhizi*–soybean pathosystem from organic and conventional soybean production systems. Plant Pathol. 67, 1697–1705. doi: 10.1111/ppa.12883

20. Koch, A., Biedenkopf, D., Furch, A., Weber, L., Rossbach, O., Abdellatef, E., Linicus, L., Johannsmeier, J., Jelonek, L., Goesmann, A., Cardoza, V., McMillan, J., Mentzel, T., and Kogel, K.H. (2016). An RNAi-based control of *Fusarium graminearum* infections through spraying of long dsRNAs involves a plant passage and is controlled by the fungal silencing machinery. PLoS Pathog. 12, e1005901. doi: 10.1371/journal.ppat.1005901

21. Kondo, T., Kose, R., Naito, H., and Kasai, W. (2014). Aqueous counter collision using paired water jets as a novel means of preparing bio-nanofibers. Carbohydrate Polymers 112, 284–290. doi: 10.1016/j.carbpol.2014.05.064

22. Kose, R., Kasai, W., and Kondo, T. (2011). Switching surface properties of substrates by coating with a cellulose nanofiber having a high adsorbability. Sen’i Gakkaishi 67, 163–168. doi: 10.2115/fiber.67.163

23. Langenbach, C., Campe, R., Beyer, S.F., Mueller, A.N., and Conrath, U. (2016). Fighting Asian soybean rust. Front. Plant Sci. 7, 797. doi: 10.3389/fpls.2016.00797

24. Lenardon, M.D., Munro, C.A., and Gow, N.A. (2010). Chitin synthesis and fungal pathogenesis. Curr. Opin. Microbiol. 13, 416–423. doi: 10.1016/j.mib.2010.05.002

25. Li, S., Smith, J.R., Ray, J.D., and Frederick, R.D. (2012). Identification of a new soybean rust resistance gene in PI 567102B. Theor. Appl. Genet. 125, 133–142. doi: 10.1007/s00122-012-1821-y

26. Madrid, M.P., Di Pietro, A., and Roncero, M.I. (2003). Class V chitin synthase determines pathogenesis in the vascular wilt fungus *Fusarium oxysporum* and mediates resistance to plant defence compounds. Mol. Microbiol. 47, 257–266. 10.1046/j.1365-2958.2003.03299.x

27. Maltby, L., Brock, T.C., and Van den Brink, P.J. (2009). Fungicide risk assessment for aquatic ecosystems: importance of interspecific variation, toxic mode of action, and exposure regime. Environ. Sci. Technol. 43, 7556–7563. doi: 10.1021/es901461c

28. McLean, R., and Byth, D. (1980). Inheritance of resistance to rust *Phakopsora pachyrhizi* in soybeans. Aust. J. Agric. Res. 31, 951–956. doi: 10.1071/AR9800951

29. Mena, E., Stewart, S., Montesano, M., and Ponce de León, I. (2019). Soybean stem canker caused by *Diaporthe caulivora*; Pathogen diversity, colonization process, and plant defense activation. Front. Plant Sci. 10, 1733.

30. Mendoza-Mendoza, A., Berndt, P., Djamei, A., Weise, C., Linne, U., Marahiel, M., Vranes, M., Kämper, J., and Kahmann, R. (2009). Physical-chemical plant-derived signals induce differentiation in *Ustilago maydis*. Mol. Microbiol. 71, 895–911.

31. Mondal, S. (2017). Preparation, properties and applications of nanocellulosic materials. Carbohydr. Polym. 163, 301–316.

32. Monteros, M.J., Missaoui, A.M., Phillips, D.V., Walker, D.R., and Boerma, H.R. (2007). Mapping and confirmation of the ‘Hyuuga’ red-brown lesion resistance gene for Asian soybean rust. Crop Sci. 47, 829–834.

33. Naoumkina, M.A., Zhao, Q., Gallego-Giraldo, L., Dai, X., Zhao, P.X., and Dixon, R.A. (2010). Genome-wide analysis of phenylpropanoid defence pathways. Mol. Plant Pathol. 11, 829–846. doi: 10.1111/j.1364-3703.2010.00648.x

34. Nerva, L., Sandrini, M., Gambino, G., and Chitarra, W. (2020). Double-stranded RNAs (dsRNAs) as a sustainable tool against gray mold (*Botrytis cinerea*) in grapevine: Effectiveness of different application methods in an open-air environment. Biomolecules 10, 200. doi: 0.3390/biom10020200

35. Schneider, K.T., van de Mortel, M., Bancroft, T.J., Braun, E., Nettleton, D., Nelson, R.T., Frederick, R.D., Baum, T.J., Graham, M.A., and Whitham, S.A. (2011). Biphasic gene expression changes elicited by *Phakopsora pachyrhizi* in soybean correlate with fungal penetration and haustoria formation. Plant Physiol. 157, 355–371. doi: 10.1104/pp.111.181149

36. Slaminko, T.L., Miles, M.R., Frederick, R.D., Bonde, M.R., and Hartman, G.L. (2008). New legume hosts of *Phakopsora pachyrhizi* based on greenhouse evaluations. Plant Dis. 92, 767–771. doi: 10.1094/PDIS-92-5-0767

37. Takeshita, N., Ohta, A., and Horiuchi, H. (2005). CsmA, a class V chitin synthase with a myosin motor-like domain, is localized through direct interaction with the actin cytoskeleton in *Aspergillus nidulans*. Mol. Biol. Cell 16, 1961–1970. doi: 10.1091/mbc.e04-09-0761

38. Treitschke, S., Doehlemann, G., Schuster, M., and Steinberg, G. (2010). The myosin motor domain of fungal chitin synthase V is dispensable for vesicle motility but required for virulence of the maize pathogen *Ustilago maydis*. Plant Cell 22, 2476–2494. doi: 10.1105/tpc.110.075028

39. Uppalapati, S.R., Ishiga, Y., Doraiswamy, V., Bedair, M., Mittal, S., Chen, J., Nakashima, J., Tang, Y., Tadege, M., Ratet, P., Chen, R., Schultheiss, H., and Mysore, K.S. (2012). Loss of abaxial leaf epicuticular wax in *Medicago truncatula irg1*/*palm1* mutants results in reduced spore differentiation of anthracnose and nonhost rust pathogens. Plant Cell 24, 353–370. doi: 10.1105/tpc.111.093104

40. Wang, M., Weiberg, A., Lin, F.M., Thomma, B.P., Huang, H.D., and Jin, H. (2016). Bidirectional cross-kingdom RNAi and fungal uptake of external RNAs confer plant protection. Nat. Plants 2, 16151. doi: 10.1038/nplants.2016.151

41. Weidenbach, D., Jansen, M., Franke, R.B., Hensel, G., Weissgerber, W., Ulferts, S., Jansen, I., Schreiber, L., Korzun, V., Pontzen, R., Kumlehn, J., Pillen, K., and Schaffrath, U. (2014). Evolutionary conserved function of barley and *Arabidopsis* 3-KETOACYL-CoA SYNTHASES in providing wax signals for germination of powdery mildew fungi. Plant Physiol. 166, 1621–1633. doi: 10.1104/pp.114.246348

42. Wytinck, N., Manchur, C.L., Li, V.H., Whyard, S., and Belmonte, M.F. (2020). dsRNA uptake in plant pests and pathogens: Insights into RNAi-based insect and fungal control technology. Plants (Basel) 9. doi: 10.3390/plants9121780

43. Yamaoka, Y., Yamanaka, N., and Akamatsu, H. (2014). Pathogenic races of soybean rust *Phakopsora pachyrhizi* collected in Tsukuba and vicinity in Ibaraki, Japan. J. Gen. Plant Pathol. 80, 184–188. doi: 10.1007/s10327-014-0507-5

44. Yorinori, J.T., Paiva, W.M., Frederick, R.D., Costamilan, L.M., Bertagnolli, P.F., Hartman, G.E., Godoy, C.V., and Nunes, J. (2005). Epidemics of soybean rust (*Phakopsora pachyrhizi*) in Brazil and Paraguay from 2001 to 2003. Plant Dis. 89, 675–677. doi: 10.1094/PD-89-0675

