## Supplemental Table for "Covering soybean leaves with cellulose nanofiber changes leaf surface hydrophobicity and confers resistance against *Phakopsora pachyrhizi*"

### Slide 1
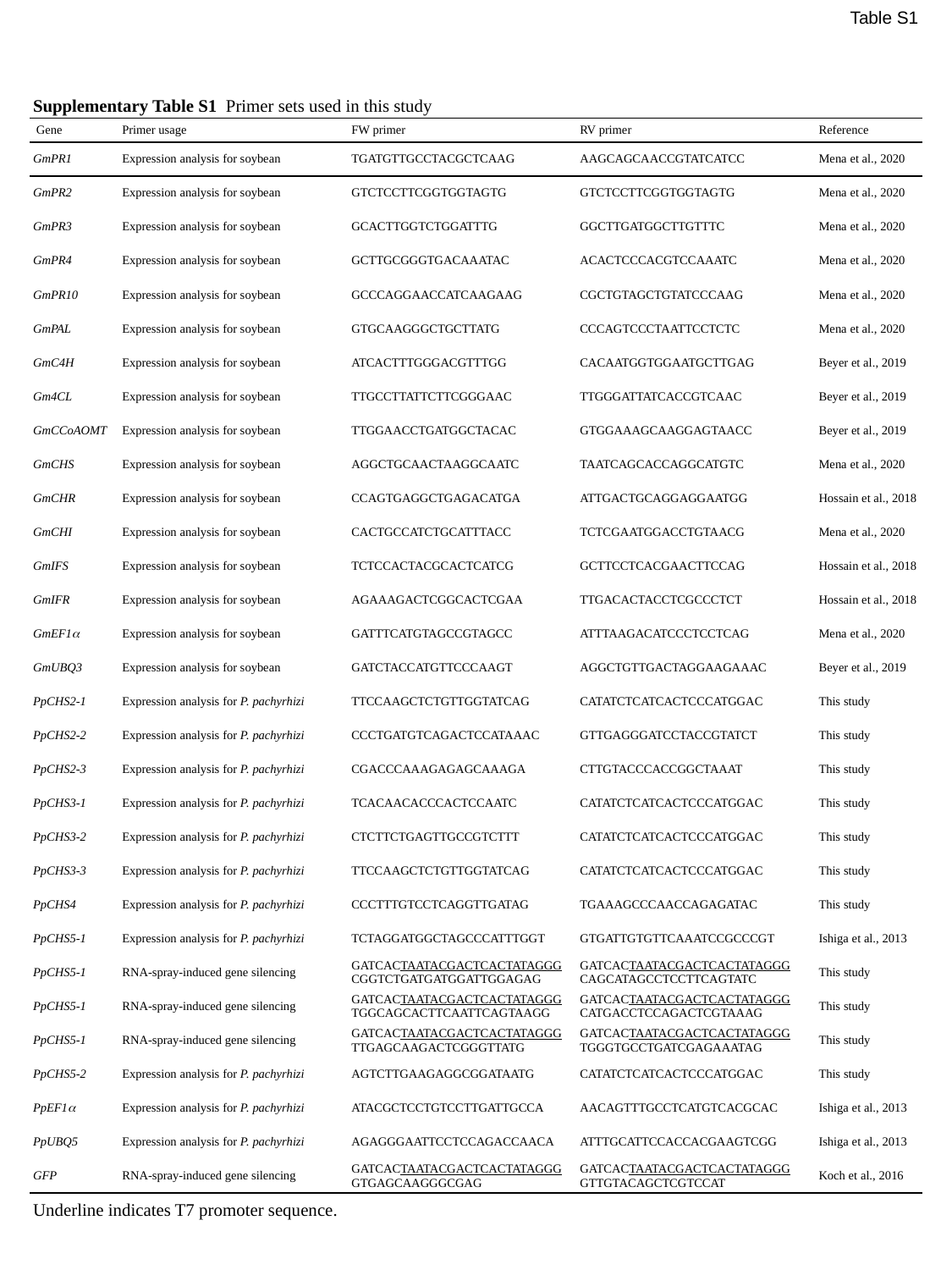

Table S1
| Supplementary Table S1 Primer sets used in this study | | | | |
| --- | --- | --- | --- | --- |
| Gene | Primer usage | FW primer | RV primer | Reference |
| GmPR1 | Expression analysis for soybean | TGATGTTGCCTACGCTCAAG | AAGCAGCAACCGTATCATCC | Mena et al., 2020 |
| GmPR2 | Expression analysis for soybean | GTCTCCTTCGGTGGTAGTG | GTCTCCTTCGGTGGTAGTG | Mena et al., 2020 |
| GmPR3 | Expression analysis for soybean | GCACTTGGTCTGGATTTG | GGCTTGATGGCTTGTTTC | Mena et al., 2020 |
| GmPR4 | Expression analysis for soybean | GCTTGCGGGTGACAAATAC | ACACTCCCACGTCCAAATC | Mena et al., 2020 |
| GmPR10 | Expression analysis for soybean | GCCCAGGAACCATCAAGAAG | CGCTGTAGCTGTATCCCAAG | Mena et al., 2020 |
| GmPAL | Expression analysis for soybean | GTGCAAGGGCTGCTTATG | CCCAGTCCCTAATTCCTCTC | Mena et al., 2020 |
| GmC4H | Expression analysis for soybean | ATCACTTTGGGACGTTTGG | CACAATGGTGGAATGCTTGAG | Beyer et al., 2019 |
| Gm4CL | Expression analysis for soybean | TTGCCTTATTCTTCGGGAAC | TTGGGATTATCACCGTCAAC | Beyer et al., 2019 |
| GmCCoAOMT | Expression analysis for soybean | TTGGAACCTGATGGCTACAC | GTGGAAAGCAAGGAGTAACC | Beyer et al., 2019 |
| GmCHS | Expression analysis for soybean | AGGCTGCAACTAAGGCAATC | TAATCAGCACCAGGCATGTC | Mena et al., 2020 |
| GmCHR | Expression analysis for soybean | CCAGTGAGGCTGAGACATGA | ATTGACTGCAGGAGGAATGG | Hossain et al., 2018 |
| GmCHI | Expression analysis for soybean | CACTGCCATCTGCATTTACC | TCTCGAATGGACCTGTAACG | Mena et al., 2020 |
| GmIFS | Expression analysis for soybean | TCTCCACTACGCACTCATCG | GCTTCCTCACGAACTTCCAG | Hossain et al., 2018 |
| GmIFR | Expression analysis for soybean | AGAAAGACTCGGCACTCGAA | TTGACACTACCTCGCCCTCT | Hossain et al., 2018 |
| GmEF1α | Expression analysis for soybean | GATTTCATGTAGCCGTAGCC | ATTTAAGACATCCCTCCTCAG | Mena et al., 2020 |
| GmUBQ3 | Expression analysis for soybean | GATCTACCATGTTCCCAAGT | AGGCTGTTGACTAGGAAGAAAC | Beyer et al., 2019 |
| PpCHS2-1 | Expression analysis for P. pachyrhizi | TTCCAAGCTCTGTTGGTATCAG | CATATCTCATCACTCCCATGGAC | This study |
| PpCHS2-2 | Expression analysis for P. pachyrhizi | CCCTGATGTCAGACTCCATAAAC | GTTGAGGGATCCTACCGTATCT | This study |
| PpCHS2-3 | Expression analysis for P. pachyrhizi | CGACCCAAAGAGAGCAAAGA | CTTGTACCCACCGGCTAAAT | This study |
| PpCHS3-1 | Expression analysis for P. pachyrhizi | TCACAACACCCACTCCAATC | CATATCTCATCACTCCCATGGAC | This study |
| PpCHS3-2 | Expression analysis for P. pachyrhizi | CTCTTCTGAGTTGCCGTCTTT | CATATCTCATCACTCCCATGGAC | This study |
| PpCHS3-3 | Expression analysis for P. pachyrhizi | TTCCAAGCTCTGTTGGTATCAG | CATATCTCATCACTCCCATGGAC | This study |
| PpCHS4 | Expression analysis for P. pachyrhizi | CCCTTTGTCCTCAGGTTGATAG | TGAAAGCCCAACCAGAGATAC | This study |
| PpCHS5-1 | Expression analysis for P. pachyrhizi | TCTAGGATGGCTAGCCCATTTGGT | GTGATTGTGTTCAAATCCGCCCGT | Ishiga et al., 2013 |
| PpCHS5-1 | RNA-spray-induced gene silencing | GATCACTAATACGACTCACTATAGGG CGGTCTGATGATGGATTGGAGAG | GATCACTAATACGACTCACTATAGGG CAGCATAGCCTCCTTCAGTATC | This study |
| PpCHS5-1 | RNA-spray-induced gene silencing | GATCACTAATACGACTCACTATAGGG TGGCAGCACTTCAATTCAGTAAGG | GATCACTAATACGACTCACTATAGGG CATGACCTCCAGACTCGTAAAG | This study |
| PpCHS5-1 | RNA-spray-induced gene silencing | GATCACTAATACGACTCACTATAGGG TTGAGCAAGACTCGGGTTATG | GATCACTAATACGACTCACTATAGGG TGGGTGCCTGATCGAGAAATAG | This study |
| PpCHS5-2 | Expression analysis for P. pachyrhizi | AGTCTTGAAGAGGCGGATAATG | CATATCTCATCACTCCCATGGAC | This study |
| PpEF1α | Expression analysis for P. pachyrhizi | ATACGCTCCTGTCCTTGATTGCCA | AACAGTTTGCCTCATGTCACGCAC | Ishiga et al., 2013 |
| PpUBQ5 | Expression analysis for P. pachyrhizi | AGAGGGAATTCCTCCAGACCAACA | ATTTGCATTCCACCACGAAGTCGG | Ishiga et al., 2013 |
| GFP | RNA-spray-induced gene silencing | GATCACTAATACGACTCACTATAGGG GTGAGCAAGGGCGAG | GATCACTAATACGACTCACTATAGGG GTTGTACAGCTCGTCCAT | Koch et al., 2016 |
| Underline indicates T7 promoter sequence. | | | | |
