## Supplemental Figures for "Covering soybean leaves with cellulose nanofiber changes leaf surface hydrophobicity and confers resistance against *Phakopsora pachyrhizi*"

### Slide 1
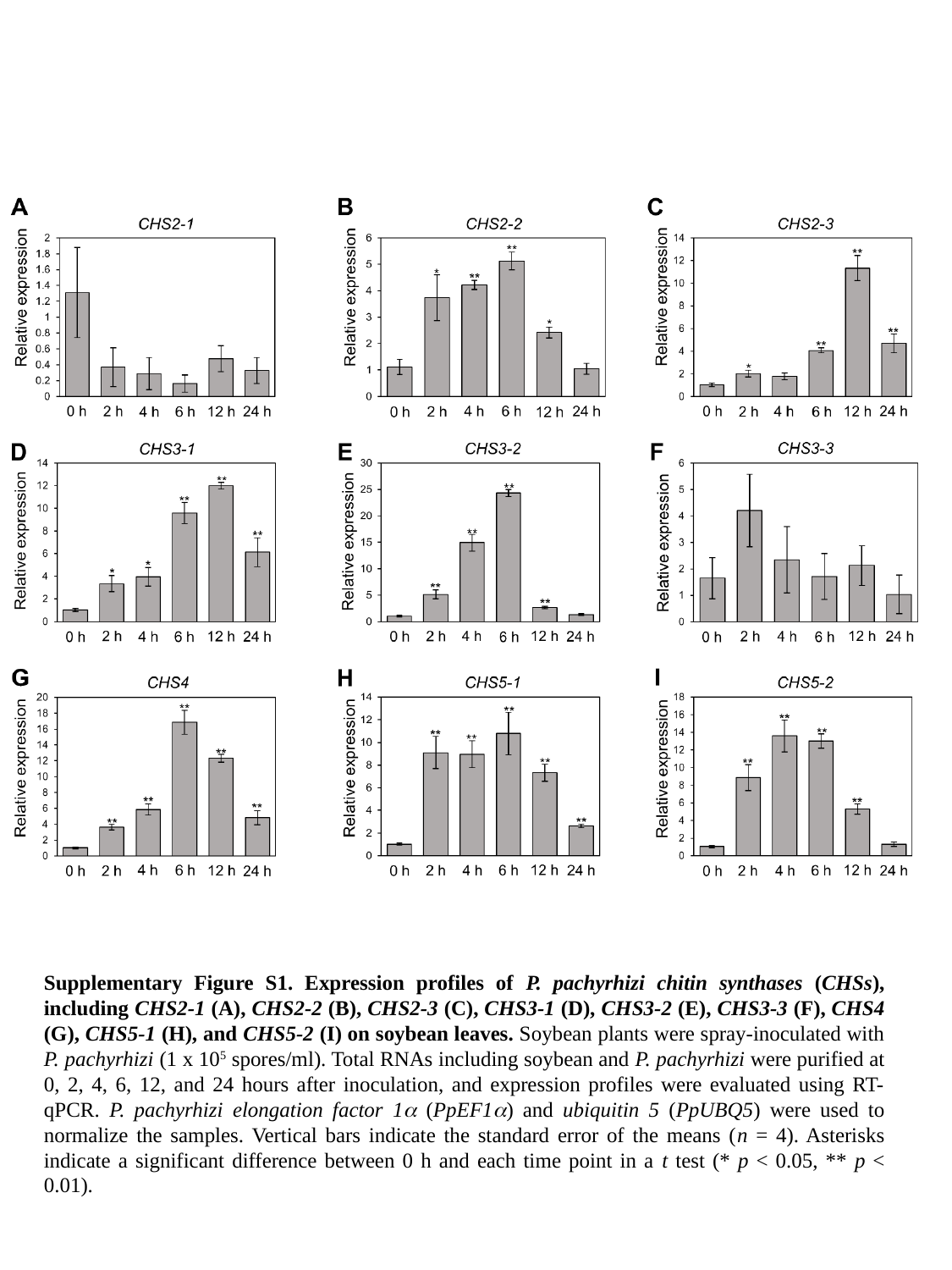

Supplementary Figure S1. Expression profiles of P. pachyrhizi chitin synthases (CHSs), including CHS2-1 (A), CHS2-2 (B), CHS2-3 (C), CHS3-1 (D), CHS3-2 (E), CHS3-3 (F), CHS4 (G), CHS5-1 (H), and CHS5-2 (I) on soybean leaves. Soybean plants were spray-inoculated with P. pachyrhizi (1 x 10^5^ spores/ml). Total RNAs including soybean and P. pachyrhizi were purified at 0, 2, 4, 6, 12, and 24 hours after inoculation, and expression profiles were evaluated using RT-qPCR. P. pachyrhizi elongation factor 1α (PpEF1α) and ubiquitin 5 (PpUBQ5) were used to normalize the samples. Vertical bars indicate the standard error of the means (n = 4). Asterisks indicate a significant difference between 0 h and each time point in a t test (* p < 0.05, ** p < 0.01).

### Slide 2
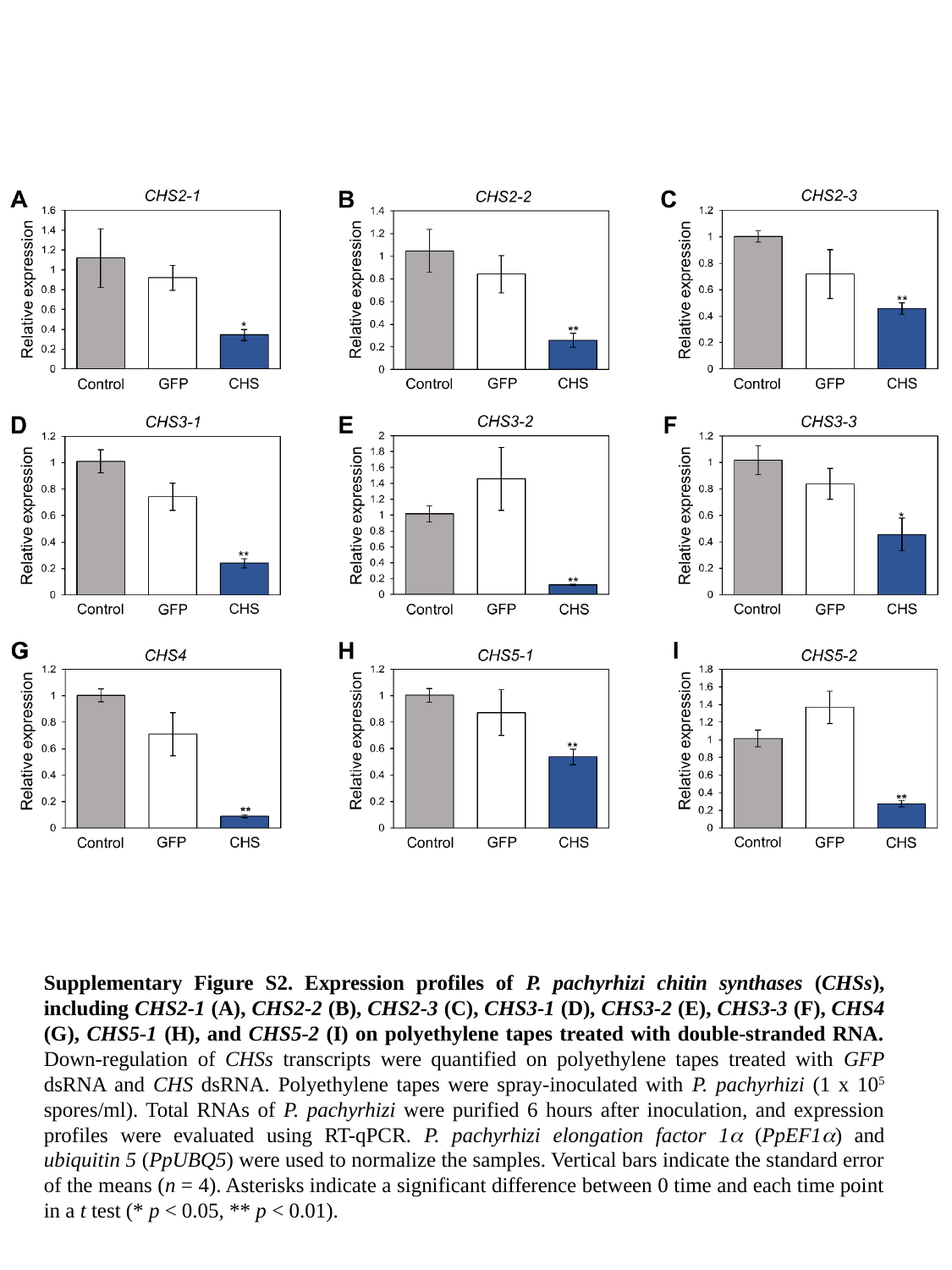

Supplementary Figure S2. Expression profiles of P. pachyrhizi chitin synthases (CHSs), including CHS2-1 (A), CHS2-2 (B), CHS2-3 (C), CHS3-1 (D), CHS3-2 (E), CHS3-3 (F), CHS4 (G), CHS5-1 (H), and CHS5-2 (I) on polyethylene tapes treated with double-stranded RNA. Down-regulation of CHSs transcripts were quantified on polyethylene tapes treated with GFP dsRNA and CHS dsRNA. Polyethylene tapes were spray-inoculated with P. pachyrhizi (1 x 10^5^ spores/ml). Total RNAs of P. pachyrhizi were purified 6 hours after inoculation, and expression profiles were evaluated using RT-qPCR. P. pachyrhizi elongation factor 1α (PpEF1α) and ubiquitin 5 (PpUBQ5) were used to normalize the samples. Vertical bars indicate the standard error of the means (n = 4). Asterisks indicate a significant difference between 0 time and each time point in a t test (* p < 0.05, ** p < 0.01).

### Slide 3
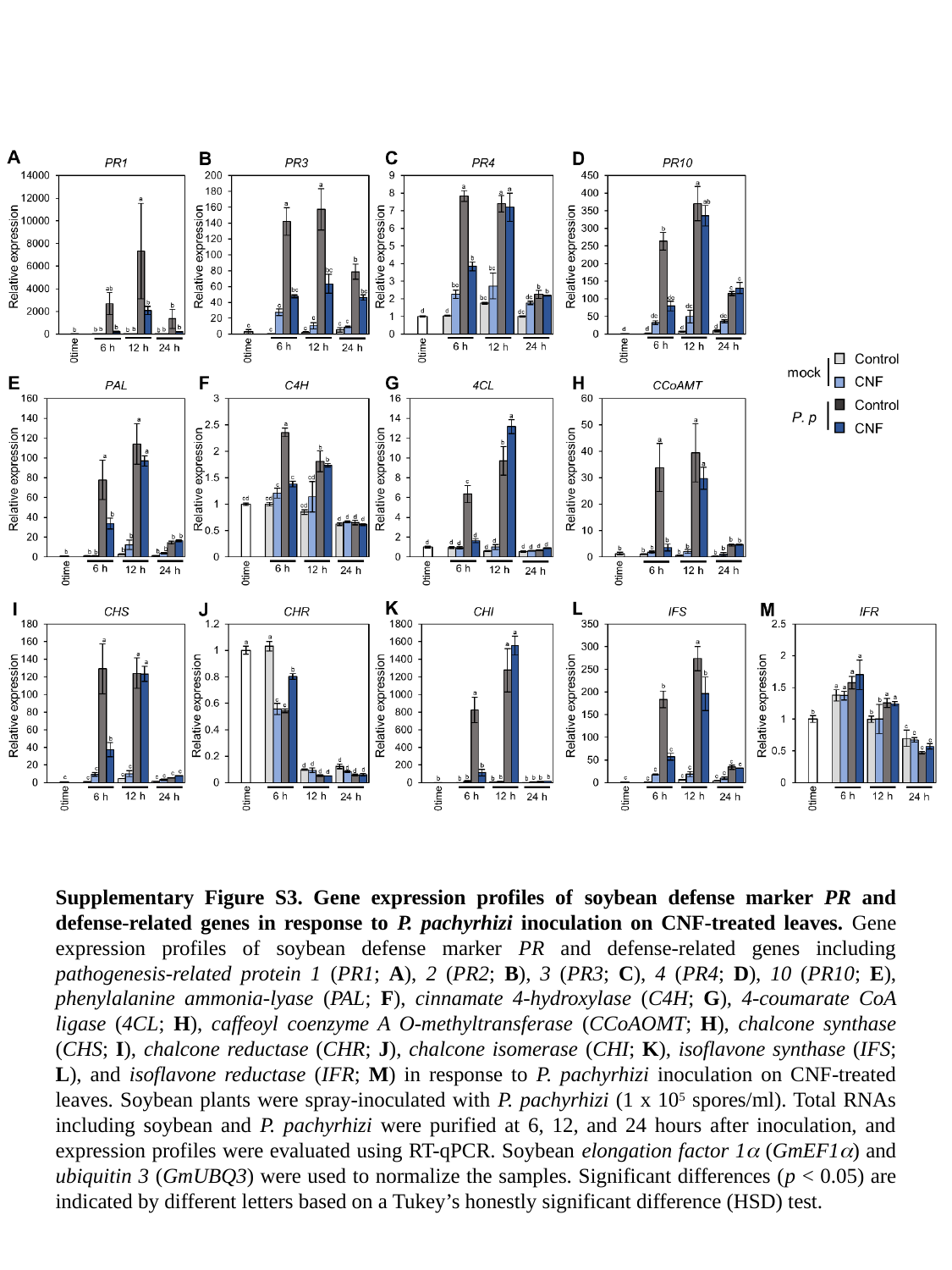

Supplementary Figure S3. Gene expression profiles of soybean defense marker PR and defense-related genes in response to P. pachyrhizi inoculation on CNF-treated leaves. Gene expression profiles of soybean defense marker PR and defense-related genes including pathogenesis-related protein 1 (PR1; A), 2 (PR2; B), 3 (PR3; C), 4 (PR4; D), 10 (PR10; E), phenylalanine ammonia-lyase (PAL; F), cinnamate 4-hydroxylase (C4H; G), 4-coumarate CoA ligase (4CL; H), caffeoyl coenzyme A O-methyltransferase (CCoAOMT; H), chalcone synthase (CHS; I), chalcone reductase (CHR; J), chalcone isomerase (CHI; K), isoflavone synthase (IFS; L), and isoflavone reductase (IFR; M) in response to P. pachyrhizi inoculation on CNF-treated leaves. Soybean plants were spray-inoculated with P. pachyrhizi (1 x 10^5^ spores/ml). Total RNAs including soybean and P. pachyrhizi were purified at 6, 12, and 24 hours after inoculation, and expression profiles were evaluated using RT-qPCR. Soybean elongation factor 1α (GmEF1α) and ubiquitin 3 (GmUBQ3) were used to normalize the samples. Significant differences (p < 0.05) are indicated by different letters based on a Tukey’s honestly significant difference (HSD) test.

### Slide 4
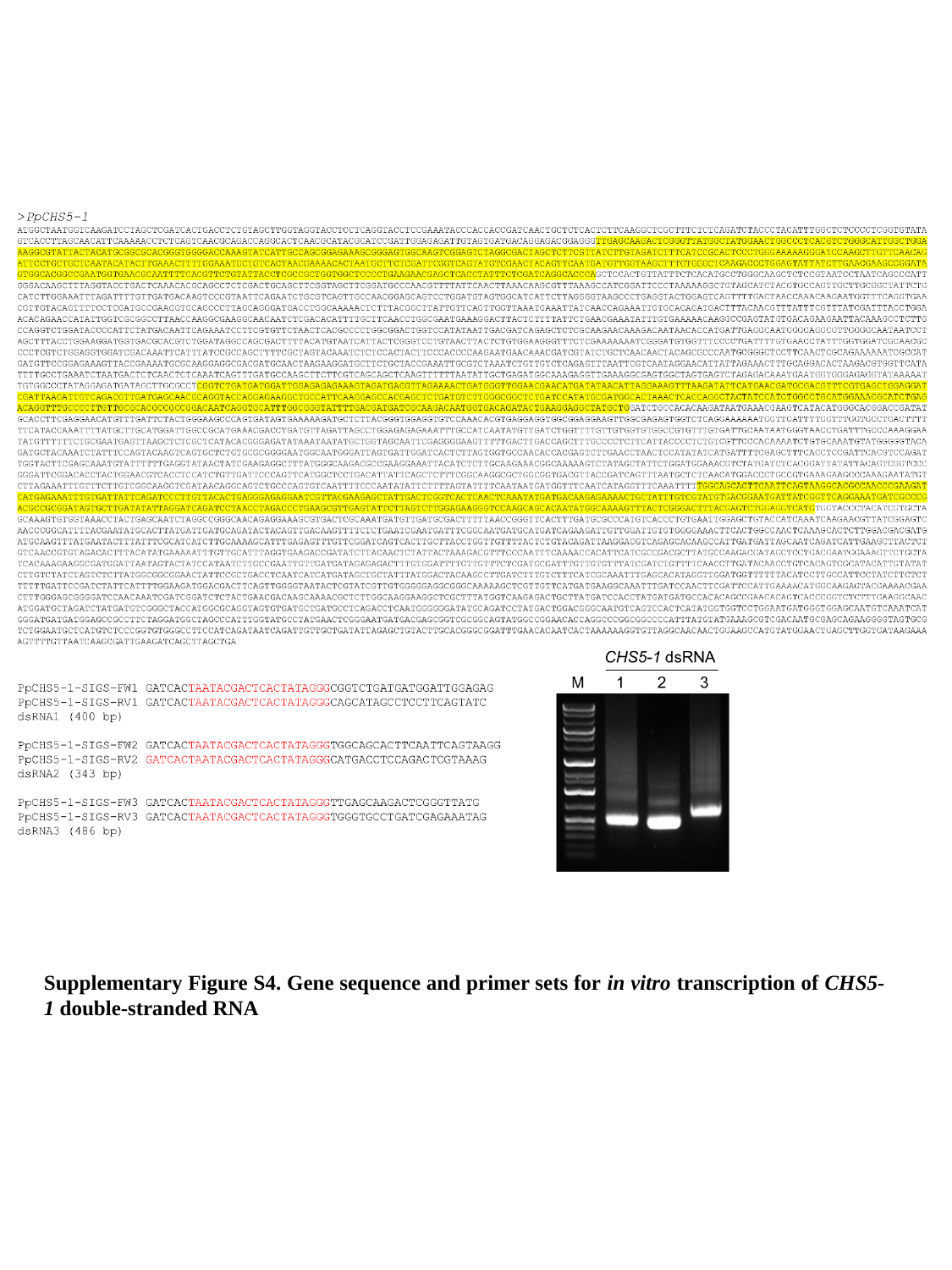

Supplementary Figure S4. Gene sequence and primer sets for in vitro transcription of CHS5-1 double-stranded RNA
